# Reshaping the progranulin/sortilin interaction for targeted degradation of extracellular proteins

**DOI:** 10.1101/2025.03.03.641244

**Authors:** Camilla Gustafsen, Joachim Vilstrup, Marianne Kristensen, Ditte Køster, Jonas Lende, Casper Larsen, Amanda Simonsen, Astrid Graversen, Ditte Olsen, Lone T. Pallesen, Christian B. Vægter, Anders Etzerodt, Omar Qureshi, Neale Harrison, Jamie Cowley, Nicholas Barnes, Sofia M. M. Mazarakis, Daniel Greve, Anna Quattropani, Paul Glossop, Gavin Whitlock, Klaus Th. Jensen, Simon F. Nielsen, Peder Madsen, Simon Glerup

## Abstract

Targeted protein degradation (TPD) using PROteolysis TArgeting Chimeras (PROTACs) is a rapidly emerging therapeutic strategy for difficult-to-drug cytosolic proteins. PROTACs are heterobifunctional small molecules that bridge the target with an E3 ubiquitin ligase, destining it for degradation by the proteasome. They have the potential to be orally available and to act catalytically, switching the pharmacology from occupancy-driven to event-driven (1-3). Here we present a strategy for targeted degradation of extracellular proteins by reshaping the interaction between the broadly expressed lysosome sorting receptor sortilin and its ligand progranulin for engineering SORtilin-based lysosome TArgeting Chimeras (SORTACs). SORTACs induce ternary complex formation with the target and sortilin, followed by endocytosis and lysosomal degradation. SORTAC activity can be genetically encoded as demonstrated by converting an IgG binding nanobody to an IgG degrading nanobody or by chemical conjugation, enabling single step conversion of therapeutic antibodies from binding their target to driving its degradation. Importantly, using structure-based design, we generated small molecule SORTACs against the inflammatory cytokine TNFa with nanomolar range potency and with physicochemical properties like PROTACs. Our results demonstrate that SORTACs constitute a versatile and highly modular platform for rapid generation of degraders of in theory any extracellular target and with the potential to have wide impact in drug discovery.

## Introduction

PROTACs are heterobifunctional small molecules consisting of an E3 ubiquitin ligase binder chemically linked to a protein target binder. In the cytosol, the PROTAC induces ternary complex formation between the target and ligase, resulting in target ubiquitylation and subsequent degradation by the ubiquitin-proteasome system (UPS). PROTACs have the potential to be orally available and can be recycled thereby acting in a sub stoichiometric manner, switching the pharmacology from occupancy-driven to event-driven. The technology also removed the need for active target binders, rendering PROTACs a particularly attractive strategy for difficult to drug targets. The PROTAC field is gaining exponential interest and several PROTACs are at advanced clinical stages in cancer and inflammatory indications. However, the PROTAC target space is exclusively limited to cytosolic proteins (3).

Extracellular proteins including soluble and transmembrane proteins constitute around 40% of the total proteome both at the level of individual gene transcripts as well as total number of transcripts (4). These encompass numerous interesting therapeutic targets that are difficult to drug with existing technologies, and include inflammatory cytokines and receptors, autoantibodies, immune complexes, protein aggregates, multidomain proteins with no druggable epitope, and scaffold proteins to name a few. Recently, technologies for targeted lysosomal degradation of extracellular proteins using antibodies or other biologics have been reported: LYTACs are based on conjugation of an antibody, a peptide or a small molecule with carbohydrate structures or *de novo* designed protein binders driving degradation via the lysosome sorting receptors CI-M6PR, ASGPR, transferrin receptor, or sortilin, respectively (5-8), SELDEGs are fusion proteins consisting of a Fc region fused to a target binding part, directing FcRn-mediated internalization and lysosomal degradation of the target (9), and KINETACs are bispecific antibodies with one arm binding to the target and the second arm consisting of the cytokine CXCL12 mediating binding to the recycling receptor CXCR7, causing lysosomal delivery of the target (10). However, all current extracellular TPD technologies rely on large or complex macromolecules, far from the physicochemical property space of PROTACs (2), and limiting their potential as broad therapeutic platforms. We here aimed to leverage sortilin, a broadly expressed lysosome sorting receptor for the secreted protein progranulin (11, 12), for developing small molecule extracellular degraders. Haploinsufficiency for progranulin causes frontotemporal dementia (FTD) and inhibiting the sortilin/progranulin interaction is safe and increases extracellular progranulin by several folds in cells, animals, and humans (13). Hence, small molecule scaffolds that bind to sortilin in the same site as progranulin have been identified (14, 15), and we hypothesized that these could serve as starting points for developing heterobifunctional molecules with similar architecture as PROTACs but targeting extracellular proteins.

## Results

### Progranulin-derived C-terminal tetrapeptide is a lysosomal sorting motif

The domain structure of progranulin contains several granulin domains organized like pearls on a string but only the C-terminal granulin E domain is required for sortilin binding (Fig. 1a) (16). We therefore speculated that fusing the progranulin C-terminal to any protein would induce sortilin binding and subsequent sortilin-mediated endocytosis. To exemplify this, we designed constructs encoding green fluorescent protein (GFP) or the inflammatory cytokine tumor necrosis factor alpha (TNFa), respectively, with a C-terminal flexible linker followed by the last four C-terminal amino acids of progranulin (RQLL) (Fig. 1b). Interestingly, the presence of the C-terminal RQLL tetrapeptide induced nanomolar affinity binding of both proteins to the purified sortilin extracellular domain (ECD) as determined using microscale thermophoresis (MST) (Fig. 1b and Extended data Fig. 1a-c). In contrast, no binding was observed in the absence of RQLL. Furthermore, C-terminal RQLL induced removal of both proteins from the culture supernatant of sortilin-expressing cells, indeed suggesting that tagging any target with the progranulin C-terminal would induce sortilin-mediated binding and internalization (Fig. 1c-d). The cardiovascular risk protein PCSK9 is a known sortilin ligand that binds sortilin with similar affinity as progranulin but unlike for progranulin, binding does not result in sortilin-mediated internalization (17). We speculated that the introduction of RQLL at the PCSK9 C-terminal could induce internalization in a sortilin-dependent manner (Fig. 1b). Indeed, only PCSK9-RQLL was internalized by sortilin expressing cells in a manner that was inhibited by the small molecule AF38469 binding in the same pocket as RQLL (Fig. 1e) (18). Thus, specifically mimicking the sortilin/progranulin interaction allows a bound ligand to be internalized by sortilin.

**Fig. 1.**
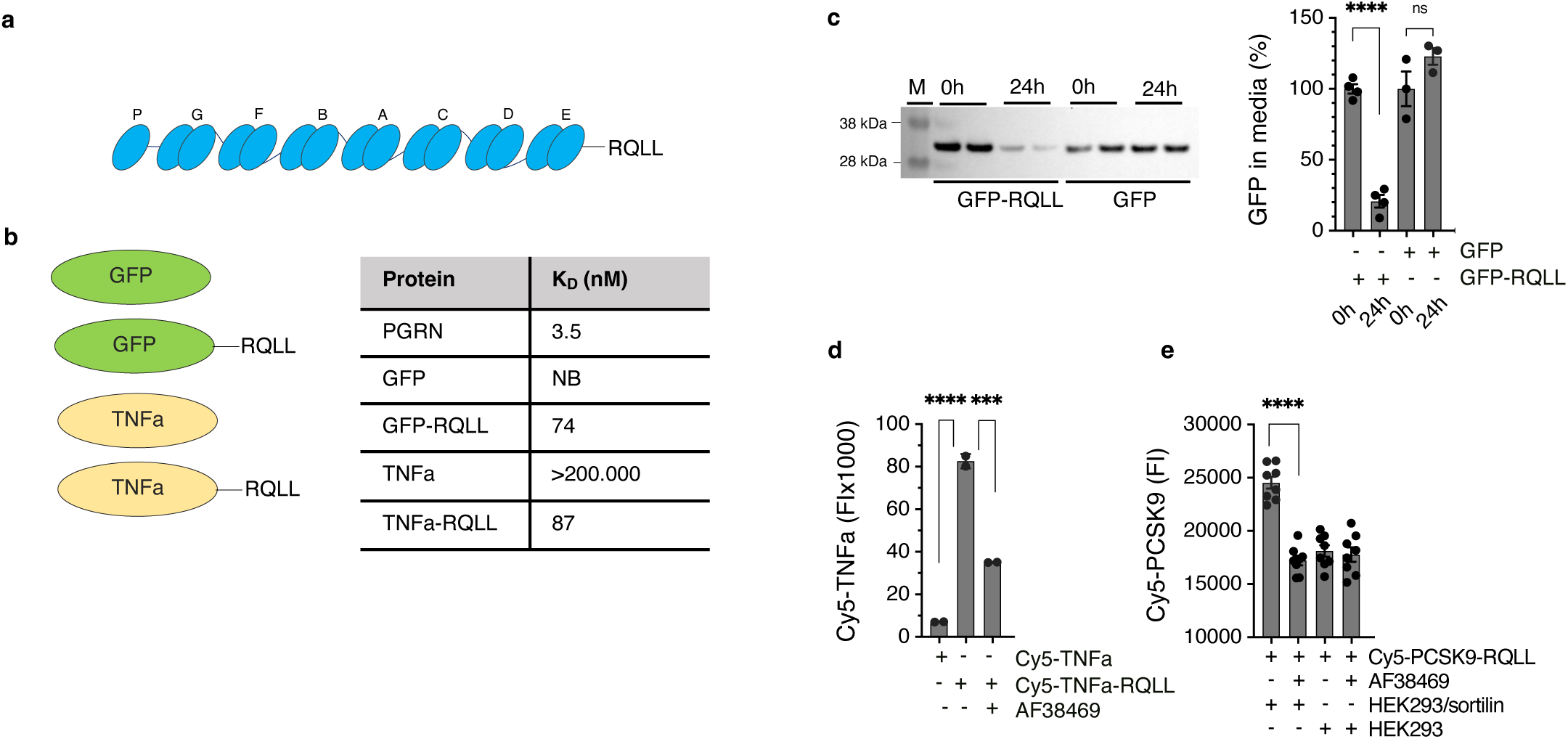
Progranulin C-terminal is a lysosome sorting tag. **a,** Model depicting progranulin domain structure indicating the individual granulin domains (P, G, F, B, A, C, D, E) with the four C-terminal amino acids RQLL. **b,** Illustration showing GFP and TNFa with or without RQLL C-terminal, conferring binding to sortilin as indicated in table. **c,** Western blot showing clearance of GFP (10 nM) with (n=4) or without (n=3) C-terminal RQLL extension from the culture supernatant of sortilin expressing HEK293 cells (HEK293/sortilin cells) following 24 h incubation. Data is normalized to GFP-RQLL at 0 h and shown as mean ± standard error of mean (SEM). **d,** Sortilin-dependent internalization (4 h) of Cy5-TNFa (100 nM) with or without C-terminal RQLL extension by HEK293/sortilin cells in the absence or presence of the small molecule sortilin inhibitor AF38469 (10 µM) (mean ± SEM, n=2). **e,** Sortilin-dependent internalization (4 h) of Cy5-PCSK9 with C-terminal RQLL extension (1 µM) in mock or sortilin expressing HEK293 cells (mean ± SEM, n=8). p-values were determined by unpaired two-tailed t-test: *p<0.05, **p<0.01, ***p<0.001, ****p<0.0001. n indicates the number of biological replicates in each experiment. Each data set shown is representative of at least three independent experiments.

### The sortilin/progranulin interaction as a basis for degrader design

Inspired by the above observations, we used RQLL as a basis for designing SORtilin-based lysosome Targeting Chimeras (SORTACs (Fig. 2a). We first conjugated RQLL to biotin to drive degradation of fluorescently labelled neutravidin (NA650) (Fig. 2b) and studied the ability of RQLL-biotin (pep-001) to drive ternary complex formation between sortilin ECD and streptavidin using a TR-FRET based assay. RQLL-biotin induced a marked increase in signal in a dose-dependent manner resulting in a bell-shaped curve with the maximum peak height at 6.3 nM (Fig. 2c), resembling the so-called hook effect of PROTACs that is caused by a concentration-dependent switch in the number of binary complexes compared to ternary complexes (1). No induction of complex formation was observed using pep-018 with an amidated C-terminal, resulting in loss of sortilin binding. We further observed that RQLL-biotin induced concentration dependent uptake of NA650 in sortilin-expressing cells, directing it to structures positive for the lysosomal marker LAMP1 (Fig. 2d) and likely leading to its degradation. This was evident by increased intact NA650 in lysates in the presence of the lysosomal proteinase inhibitor leupeptin (Fig. 2e). Uptake was completely inhibited by excess isolated biotin, RQLL peptide (pep-010), or the small molecule sortilin binder AF38469 (18) (Fig. 2f-h), showing that the effect was specifically dependent on intact binding to both sortilin and NA650. SORTAC activity was further confirmed across a range of cell lines with endogenous sortilin expression (Extended data Fig. 2a). Notably, we observed that a single addition of RQLL-biotin to NA650 containing culture medium of sortilin expressing cells led induced a cumulative depletion of NA650 over several days (Fig. 2i), suggesting that the SORTAC mechanism operates in sustained manner and is not desensitized over time.

**Fig. 2.**
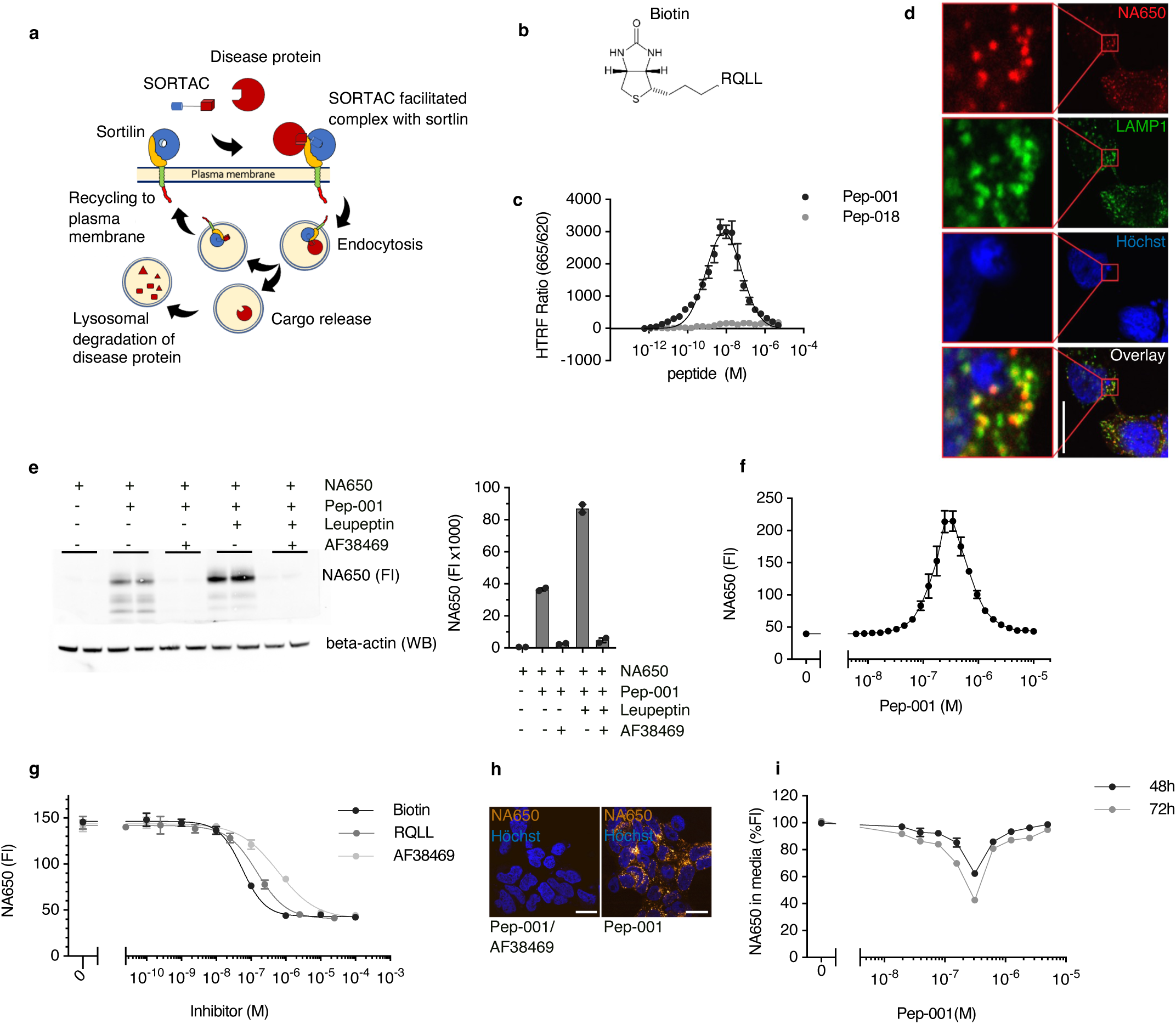
The progranulin C-terminal as a basis for degrader design. **a,** Depiction of the mechanism of action of SORtilin-based lysosome TArgeting Chimeras (SORTACs) for driving degradation of extracellular targets in lysosomes. **b,** Illustration of a SORTAC based on RQLL conjugated to biotin. **c,** SORTAC-induced (pep-001) ternary complex formation between sortilin ECD and streptavidin as measured using TR-FRET. An inactive SORTAC (pep-018) with amidated C-terminal, disrupting sortilin binding was included as negative control. Data is shown as mean ± SEM (n=3). **d,** Confocal microscopy showing SORTAC-induced (2 µM pep-001) internalization (4 h) of NA650 (500 nM) in HEK293/sortilin cells, directing it to structures positive for the lysosomal marker LAMP1. Scale bar, 20 µm. **e,** Western blot showing sortilin-dependent SORTAC-induced (300 nM pep-001) NA650 (100 nM) internalization and degradation in HEK293/sortilin cells following 24 h incubation in the absence or presence of the lysosomal proteinase inhibitor leupeptin (80 µM), and in the absence or presence of the sortilin inhibitor AF38469 (10 µM). Beta-actin was included as loading control. Quantification of the NA650 fluorescence intensity (FI) signal is shown as bar graphs (mean ± SEM, n=2). **f,** Dose-dependent pep-001 induced internalization (4 h) of NA650 (100 nM) in HEK293/sortilin cells. Data shown as mean ± SD (n=4). **g-h,** SORTAC-induced (300 nM pep-001) internalization (4 h) of NA650 (100 nM) in HEK293/sortilin cells was blocked in a dose-dependent manner by excess isolated biotin, isolated RQLL peptide (pep-010), or AF38469. Data is shown as mean ± SD (n=4). **i,** A single addition of SORTAC to the culture medium of HEK293/sortilin cells induces a sustained depletion of NA650 (100 nM) over time in a dose-dependent manner. Data shown as mean ± SEM (n=3). n indicates the number of biological replicates in a given experiment. Each data set shown is representative of at least three independent experiments.

### Genetic encoding of degrader activity in single domain antibody targeting IgG

To understand if the sortilin/progranulin interaction could shape the design of targeted degraders of clinically relevant extracellular proteins, we genetically engineered a single domain antibody (V_H_H fragment/nanobody) binding to kappa lights chains (LC) of IgG (19) to contain an extended RQLL C-terminal (Nb-RQLL) (Fig. 3a). Targeted degradation of IgG and specifically light chains is potentially relevant in several indications, including autoimmune disease (20), light-chain deposition disease (21), and light-chain amyloidosis (22). The resulting SORTAC nanobody maintained IgG binding as demonstrated using gel filtration chromatography (Extended data Fig. 3a), and induced dose-dependent internalization (Fig. 3b-c) and complete removal of IgG from the culture medium over the course of 24 hours (Fig. 3d). Sortilin levels remained constant during the experiment while no IgG accumulation was observed in lysates, suggesting that IgG gets degraded following SORTAC-induced internalization. Indeed, inhibition of lysosomal proteolysis by the presence of leupeptin increased intact IgG heavy and light chains (HC and LC, respectively) in lysates (Fig. 3e). Taken together, these findings illustrate a strategy for the swift genetic conversion of a protein-based target binder to a degrader.

**Fig. 3.**
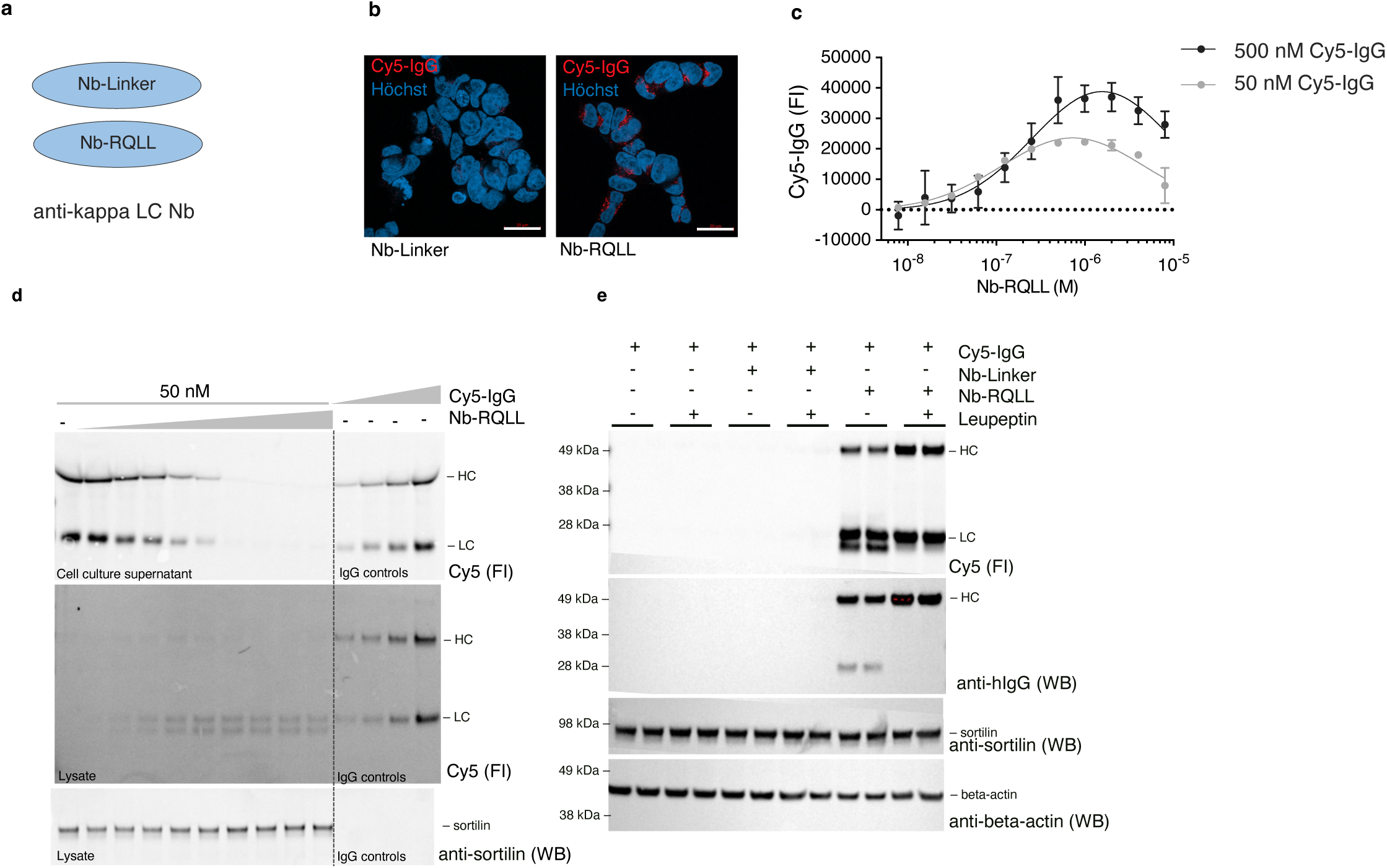
Genetically encoded SORTAC targeting IgG. **a,** Depiction of a nanobody directed against IgG kappa light chains (LC), and consisting of a single domain antibody with or without an extended RQLL C-terminal (Nb-RQLL or Nb-Linker, respectively). **b,** Nb-RQLL but not Nb-Linker (250 nM) induces internalization of Cy5-labelled IgG (50 nM) (red) by HEK293/sortilin cells. Nuclei are stained with Hoechst (blue). Scale bars, 20 µM. **c,** Nb-RQLL SORTAC-induced dose-dependent internalization of Cy5-IgG (50 nM or 500 nM) (mean ± SEM, n=2). **d,** Western blot of Cy5 fluorescent intensity (Cy5 FI) showing dose-dependent Nb-RQLL SORTAC (7.8 -2000 nM) removal of Cy5-IgG (50 nM) from the culture supernatant of HEK293/sortilin cells over the course of 24 h. The four lanes to the right contain unconditioned medium containing IgG (6.25, 12.5, 25, 50 nM IgG) as controls. The middle panel shows that IgG HC and LC do not accumulate in the cell lysates. The lower panel Western blot with anti-sortilin antibody shows that sortilin levels remains constant independent of increasing SORTAC concentration. Ig heavy chain (HC), light chain (LC) and degradation products indicated. **e,** On-blot fluorescence (Cy5) and Western blot analysis showing that Nb-RQLL (250 nM) but not Nb-Linker induces internalization of Cy5-IgG (50 nM). IgG is subsequently degraded as indicated by increased band intensity in the presence of leupeptin (80 µM). Western blot of sortilin and beta-actin were included as controls. Ig heavy chain (HC) and light chain (LC) are indicated. n indicates the number of biological replicates in each experiment. Each data set shown is representative of at least three independent experiments.

### Single step conversion of mAbs to degraders

To enable rapid conversion of any target-binding protein to a SORTAC, we developed a chemical conjugation tool based on an NHS ester group fused to a GCG flexible linker followed by RQLL (Fig. 4a), enabling conjugation of the sortilin binding peptide to primary amines on the target binder. We first conjugated the therapeutic PCSK9 inhibitory antibody alirocumab with NHS-RQLL or Linker-NHS as control. Intact binding of PCSK9 was confirmed for both alirocumab-RQLL (Ali-RQLL) and alirocumab-linker (Ali-linker) by gel filtration, showing that PCKS9 and antibody eluted as a complex (Extended Data Fig. 4a-b). Ali-RQLL but not Ali-Linker induced concentration-dependent ternary complex formation with PCSK9 and sortilin ECD as assessed by TR-FRET with a peak value of 0.07 nM (Fig. 4b). Similarly, only Ali-RQLL induced cellular uptake of Cy5-labelled PCSK9 in sortilin expressing cells in a bell-shaped manner (Fig. 4c). We next speculated that conjugation with the progranulin C-terminal RQLL could be a general strategy for converting a therapeutic mAb to a degrader in a single step reaction. Indeed, conjugation of the therapeutic TNFa inhibitory mAb adalimumab to RQLL (Ada-RQLL) conferred binding to sortilin ECD (Extended Data Fig. 4c), induced ternary complex formation with TNFa and sortilin (Fig. 4d), and dose-dependent internalization of TNFa in sortilin expressing cells as well as depletion of extracellular TNFa from the culture supernatant (Fig. 4e-f). No effect was observed for control conjugated adalimumab (Ada-linker). Of note, we observed that sortilin also mediated internalization of Ada-RQLL alone but not Ada-Linker (Fig. 4f and Extended Data Fig. 1c). Similarly, RQLL conjugation of the therapeutic mAb targeting the inflammatory cytokine IL-17A secukinumab (Sec-RQLL) conferred binding to sortilin ECD (Extended Data Fig. 4d), induced ternary complex formation with IL-17A and sortilin, and IL-17A internalization (Fig. 4g).

**Fig. 4.**
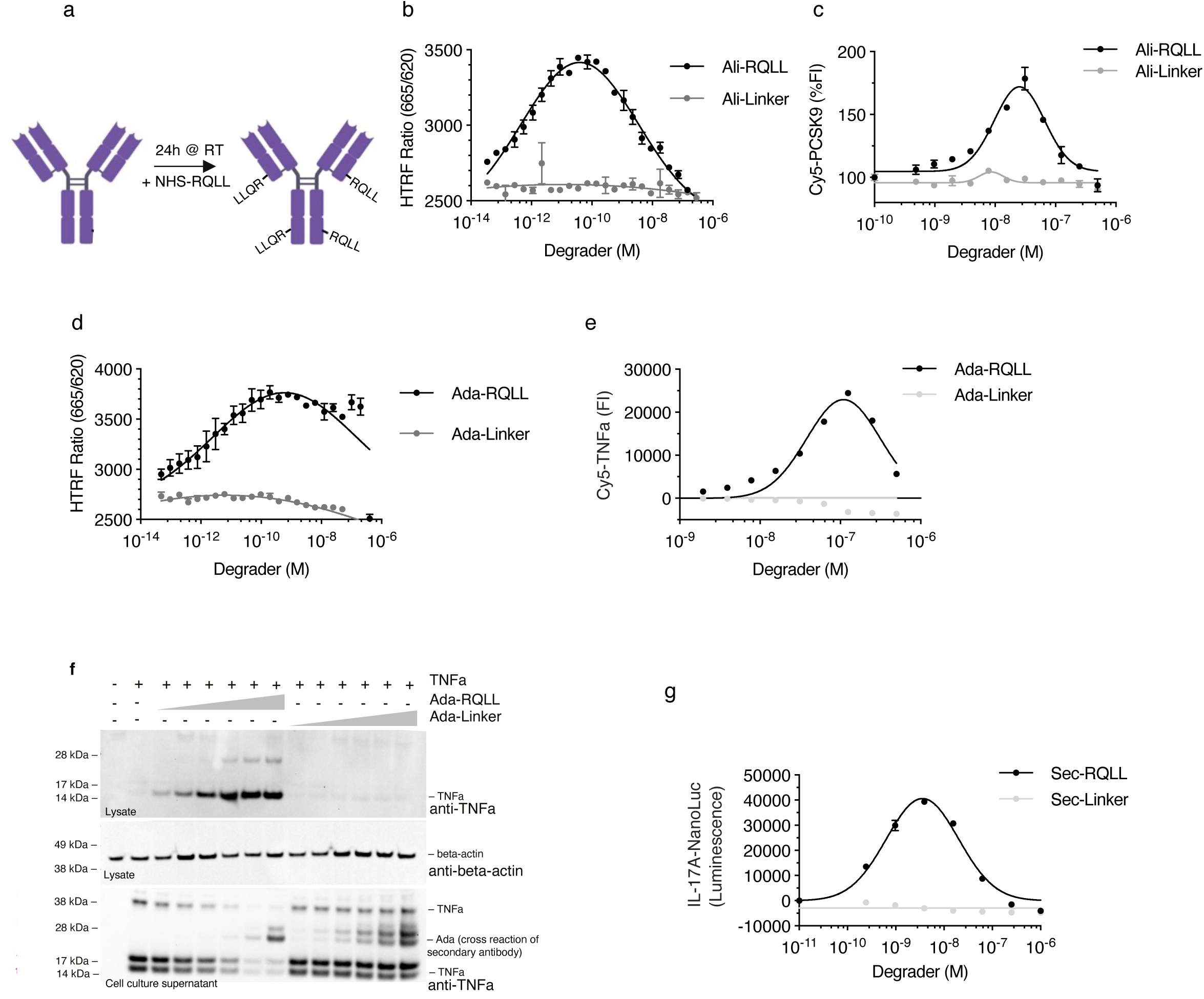
Single-step conversion of therapeutics mAbs to SORTACs. **a,** Strategy for single step conversion of a mAb to a SORTAC using a reactive NHS ester chemical tool fused to a flexible linker followed by RQLL (NHS-RQLL). **b,** TR-FRET demonstrating ternary complex formation between PCSK9 and sortilin ECD induced by alirocumab converted to a SORTAC (Ali-RQLL) in a concentration dependent manner. Control conjugated alirocumab (Ali-Linker) did not induce ternary complex formation. Data shown as mean ± SD (n=2). **c,** Ali-RQLL but not Ali-Linker induces increased internalization of Cy5-PCSK9 (100 nM) by HEK293/sortilin cells in a concentration-dependent manner. Data shown as mean ± SEM (n=2). **d,** TR-FRET demonstrating ternary complex formation between TNFa and sortilin ECD induced by adalimumab converted to a SORTAC (Ada-RQLL) but not by control conjugated adalimumab (Ada-Linker). Data shown as mean ± SEM (n=3). **e,** Dose-dependent internalization of Cy5-TNFa (100 nM) induced by Ada-RQLL but not Ada-Linker in HEK293/sortilin cells. Data shown as mean ± SEM (n=2). **f,** Western blot showing dose-dependent cellular uptake of TNFa (100 nM) by Ada-RQLL but not Ada-Linker (0.1-10 µM) (upper panel). Similarly, dose-dependent removal of TNFa from the culture supernatant was induced by Ada-RQLL but not the control Ada-Linker (lower panel). Western blot for beta-actin in lysates is shown as loading control (middle panel). Some cross reaction of the secondary antibody with adalimumab is observed (double band marked Ada). **g,** RQLL conjugation of the therapeutic mAb targeting the inflammatory cytokine IL-17A secukinumab (Sec-RQLL) induces dose-dependent internalization of NanoLuc®-IL-17 A by HEK293/sortilin cells following 3 hours incubation. No induction of internalization was observed using control conjugated secukinumab (Sec-Linker). Data is shown as mean ± SEM (n=2). n indicates the number of biological replicates in each experiment. Each data set shown is representative of at least three independent experiments.

### Design of small molecule extracellular degraders

Designing small molecule SORTAC followed the same principles as for mAbs: a sortilin binder and a warhead for the protein of interest connected via a linker. Small molecule binders inhibiting the sortilin/progranulin interaction and crystal structures of their complexes with sortilin have been reported (15, 18). Thus, inspection of the published sortilin/small molecule binder crystal structures allowed us to identify positions for exit vectors on sortilin binders without compromising binding to sortilin (Fig. 5a). This was confirmed by synthesizing and testing several sortilin binders with small PEG substituents in various positions. The best of these were used in our first proof-of-concept studies (Fig. 5a). We connected the sortilin binder via simple PEG linkers of variable length to a biotin warhead (Fig. 5b) and tested their ability to induce ternary complex formation with sortilin ECD and streptavidin by TR-FRET (Fig. 5c). The TR-FRET signal increased with increasing linker length, likely because the biotin binding site is buried relatively deep in the streptavidin quaternary structure (23, 24). Bifunctional small molecule (BSM) biotin-SORTACs (BSM1-4) induced dose-dependent cellular uptake and depletion of NA650 from the culture supernatant of sortilin expressing HEK293 cells (Fig. 5d-e). We confirmed the activity of the small molecule SORTAC mechanism in primary mouse cortical neurons (Fig. 5f) and rat Schwann cells (Fig. 5g). Importantly, reverting the stereochemistry of the sortilin binder gives compounds not binding to sortilin but with identical physicochemical properties (15). We used this to demonstrate that SORTAC activity was dependent on sortilin binding as replacing the sortilin binding part with an inactive enantiomer resulted in complete loss of induced uptake as well as affinity for sortilin (Extended Data Fig. 5a-b).

**Fig. 5.**
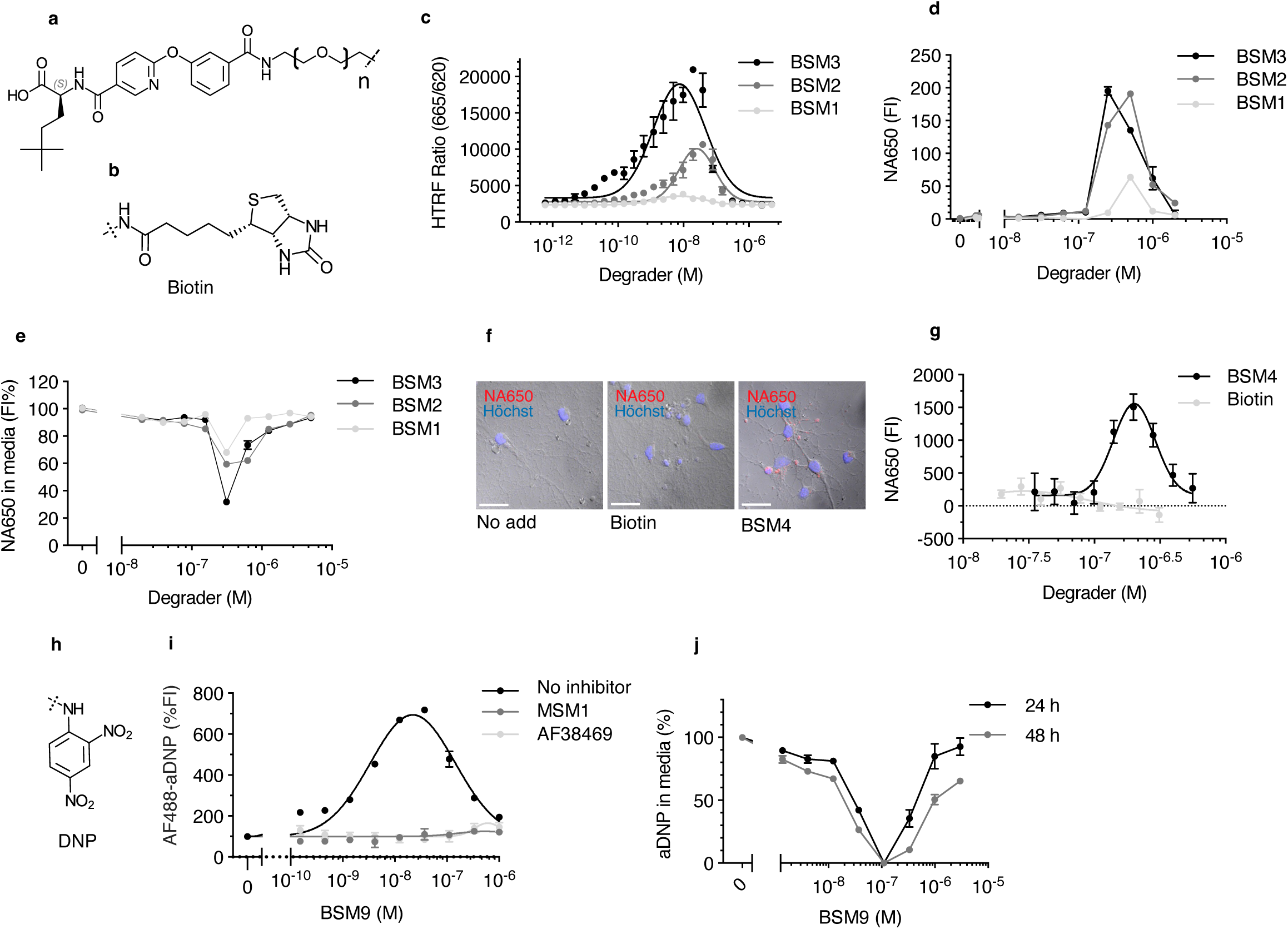
Generation of small molecule SORTACs. **a-b,** Design of small molecule SORTACs using biotin as target binder. **c-e,** Three examples of small molecule SORTACs (BSM1-3) with varying linker composition inducing ternary complex formation between sortilin ECD and streptavidin as measured by TR-FRET (mean ± SEM, n=3) (**c**), as well as internalization of NA650 (100 nM, 24 h) by HEK293/sortilin cells (mean±SD, n=2) (**d**), and clearance of NA650 (100 nM, 72 h) from the culture supernatant (mean ± SD, n=2) (**e**) in a dose-dependent manner (SORTAC concentration range 20 nM - 5 µM). **f,** Small molecule SORTAC BMS4 (624 nM) induces internalization of NA650 (100 nM, red) in primary mouse cortical neurons (DIV 13). No internalization is observed in control neurons incubated without SORTAC or with biotin alone (624 nM). Nuclei are visualized using Hoechst (blue). 24 h incubation. Scale bars are 20 µm. **g,** SORTAC activity in rat primary Schwann cells, following 24 h incubation with NA650 (100 nM) and the small molecule SORTAC BSM4 or biotin alone (20-400 nM). **h,** SORTACs targeting IgG were designed with 2,4-dinitrophenol (DNP) as a model small molecule IgG-binding antigen. **i,** SORTAC (BSM9) (2 nM - 1 µM) induced dose-dependent cellular uptake of Alexa Fluor 488 labelled rabbit polyclonal anti-DNP IgG (aDNP-AF488) (100 nM) following 24 h incubation in K562 cells, an effect that was inhibited by excess isolated sortilin monofunctional small molecule (MSM) binders MSM1 and AF38469 (10 µM), respetively. Data shown is as mean ± SEM (n=2). **j,** Rat anti DNP mAb (100 nM) internalization by HEK293/sortilin cells was induced by BSM9 in a dose- and time-dependent manner, resulting in complete removal from the culture supernatant. Data shown as mean ± SD (n=2). n indicates the number of biological replicates in each experiment. Each data set shown is representative of at least three independent experiments.

To prove the modularity of the approach, we next explored the possibility of developing small molecule degraders of IgG and applied the pesticide 2,4-dinitrophenol (DNP) as a model small molecule IgG-binding antigen (Fig. 5h). As expected, DNP-SORTACs (BSM7-9) induced dose-dependent cellular uptake of fluorescently labelled rabbit polyclonal anti-DNP IgG in both K562 cells and sortilin expressing HEK293 cells, an effect that was inhibited by excess isolated sortilin binders (Fig. 5i). Importantly, cellular fluorescence increased in the presence of leupeptin, suggesting that internalized IgG was degraded in lysosomes (Extended Data Fig. 5d). Similarly, a rat anti-DNP mAb was internalized in a dose-dependent manner and completely removed from the culture supernatant over the course of 24 hours (Fig. 5j). No effect was observed with a SORTAC based on the inactive enantiomer of the sortilin binder (BSM10), showing the critical requirement of intact sortilin binding (Extended Data Fig. 5f-g).

### Small molecule degraders of TNFa

Cytokines are an attractive but challenging class of drug targets and so far, no small molecule cytokine inhibitors have been approved. High affinity small molecule binders for TNFa have been identified but these fail to reach the efficacy of TNFa inhibitory antibodies (25). Based on one of the published TNFa/small molecule crystal structures (26), we designed and synthesized TNFa targeting SORTACs (Fig. 6a). To show the modularity of the concept, we combined the TNFa binder via linkers to 3 different sortilin binders. These induced dose-dependent cellular uptake of Cy5-TNFa (100 nM) and extracellular depletion with observed internalization already in the nanomolar range (Fig. 6b-c). The uptake was inhibited by the presence of excess isolated sortilin binder or adalimumab, suggesting that SORTAC activity was specifically dependent on engagement of both sortilin and TNFa (Extended Data Fig. 6a-b). Similarly, a SORTAC (BSM16) based on the inactive enantiomer of the sortilin binder did not induce cellular uptake (Extended Data Fig. 6c). Internalized TNFa appeared as three bands as observed by Western blotting following reducing SDS-PAGE and rapidly disappeared over time when the culture supernatant was replaced for medium without SORTAC and TNFa (Extended data Fig. 6d). We further performed uptake experiments using Cy5-TNFa in the presence or absence of lysosomal proteinase inhibition and observed that the Cy5 signal markedly increased in the presence of leupeptin (20 µM), suggesting SORTAC-directed lysosomal degradation of TNFa (Extended data Fig. 6e). This is supported by time-dependent signal observed from internalization of TNFα conjugated with a pH-sensitive fluorophore (pHrodo-TNFa), which emits light upon internalization as the pH decreases in the endosomal-lysosomal system (Extended data Fig. 6f). These findings were confirmed in human macrophages (Fig. 6d) and rat Schwann cells showing potent SORTAC-induced TNFa uptake and degradation. No uptake was observed in Schwann cells from sortilin knockout rats, confirming the specificity of the degrader mechanism (Fig. 6e and Extended Data Fig. 6f-g). Taken together, we have shown the SORTAC modularity of having different sortilin binders PEG-linked to different target warheads.

**Fig. 6.**
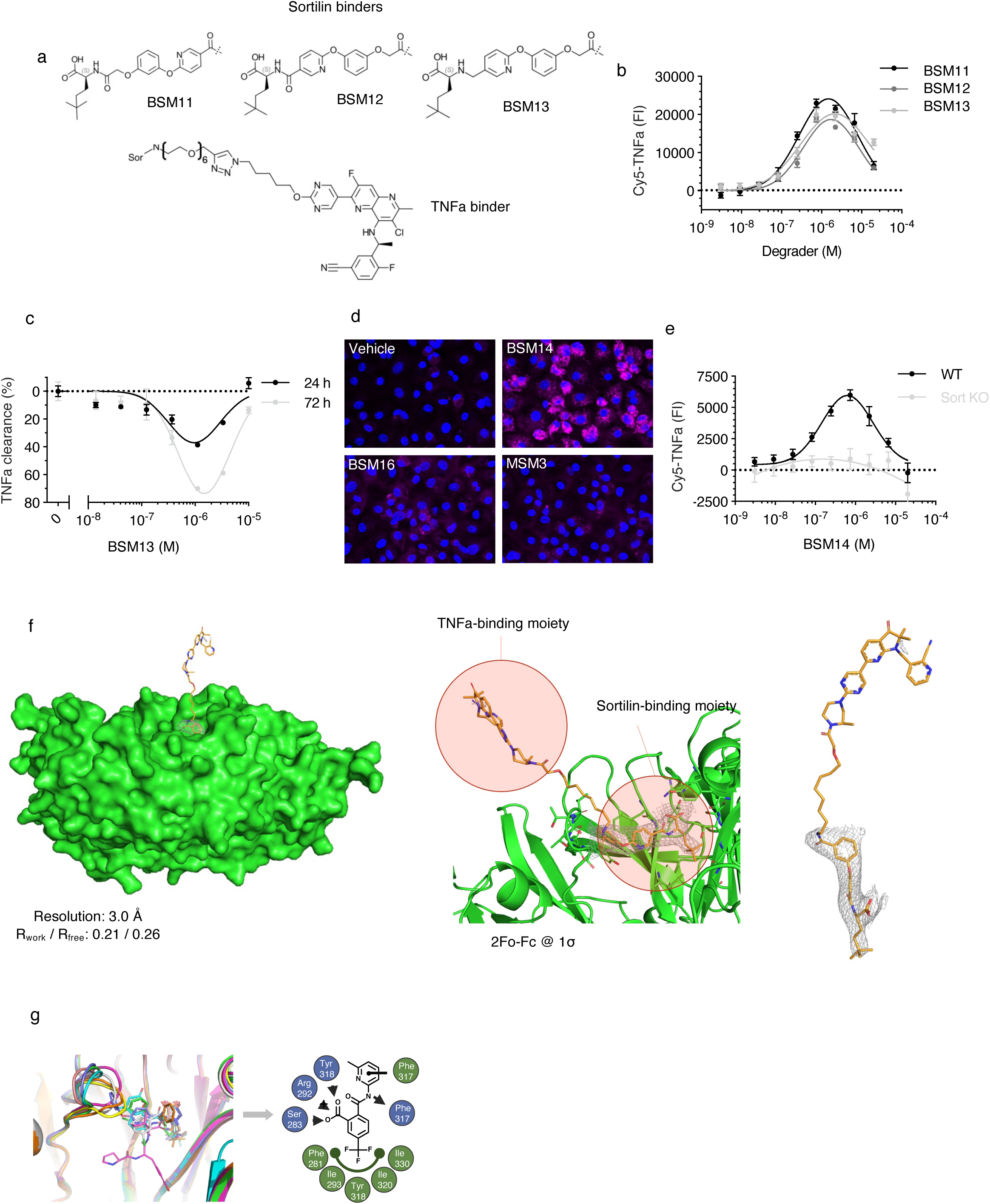
Small molecule SORTACs targeting TNFa. **a,** Small molecule TNFa targeting SORTACs were designed and synthesized with three different sortilin binding motifs. **b-c,** SORTACs (2 nM -20 µM) (BSM11-13) induced dose-dependent cellular uptake of Cy5-TNFa (100 nM) and extracellular depletion of TNFa (20 nM) in HEK293/sortilin cells. Mean value ± SEM, n=2. **d-e,** SORTAC induced TNFa uptake was also observed in macrophages derived from human monocytes (4 h incubation) (**d**) and rat Schwann cells (24 h incubation) (mean +/-SEM, n=2) (**e**) using 100 nM Cy5-TNFa and optimized SORTAC BSM14 (6.3 µM or as indicated). No induction of TNFa internalization was observed with the sortilin monobinder MSM3 (6.3 µM), the sortilin non-binding control SORTAC BSM16 (6.3 µM) or sortilin KO Schwann cells (Sort KO). **f,** Representation of the crystal structure of sortilin (left: surface view in green, middle: cartoon in green) in complex with SORTAC (sticks in orange) (BSM15). The sortilin- and TNFa-binding moiety separated by a linker is highlighted (middle). 3.0 Å resolution 2F_o_-F_c_ density map of SORTAC contoured at 1.0 s is shown (middle, right). Weak/no electron density around linker and TNFa-binding moiety indicates large flexibility in this region. **g,** Residue mapping of the sortilin hydrophobic pocket reveals binding site of RQLL tetrapeptide and small molecule ligands and highlights important interactions, guiding future small molecule SORTAC design. n indicates the number of biological replicates in each experiment. Each data set shown is representative of at least three independent experiments.

### Structural basis of SORTAC/sortilin interaction

To further elucidate the structural basis for SORTAC activity, we co-crystallized sortilin ECD with different TNFa SORTACs. Electron density is observed for sortilin ECD and the sortilin binding part of SORTACs, allowing for analysis of the binding mode between these two moieties (Fig. 6g). The binding site is located at the edge of blade 6 in the 10-bladed b-propeller domain cavity (binding site 1), as reported for known small molecule sortilin binders (15, 18). The interaction shows features like the binding mode of neurotensin, and antagonists derived from this small peptide (14, 27). Most notably is salt bridge formation to Arg292 and several interactions in the nearby hydrophobic pocket (Fig. 6g). In this case, Phe281, Ile294, Tyr318, Ile320 and Met330 contribute with hydrophobic interactions to the trimethyl-part of the SORTACs. Additional interactions involve p-stacking to the sidechain of Phe317 and hydrogen-bonds to sidechains of Tyr362, Ser319 and Ser283 or back-bone of Tyr318. No density is observed for the linker-/exit vector- and TNFa binding part, likely indicating significant flexibility in this region. The SORTAC/sortilin crystal structure demonstrates the specificity of the interaction and that it occurs at the binding site identical to that of progranulin (28), and enabling future design of optimized degraders.

## Discussion

The lysosome is the main degradative organelle of the cell with an eminent capacity for breakdown of large biomolecules and complexes, mediated by a battery of different enzymes, providing an attractive opportunity for targeted degradation of disease associated proteins (29). During biosynthesis, lysosomal enzymes are tagged with mannose-6-phosphate in the trans-Golgi network (TGN) allowing recognition by CI-M6PR and subsequently sorted to their destination in lysosomes (30). Specific substrates for lysosomal degradation are taken up from the extracellular space via receptor-mediated endocytosis. Prominent examples of lysosome sorting receptors are the low-density lipoprotein (LDL) receptor (LDLR), responsible for clearance of LDL particles by the liver (31), and the related receptor LRP2/megalin, responsible for albumin reabsorption in the kidney proximal tubule (32). LDLR and LRP2 capture their cargo at cell surface, followed by constitutive and ligand-independent endocytosis via clathrin-coated pits. The cargo is dropped at the lower pH in late endosomes, also inducing dimerization of the receptor and recycling back to the cell surface whereas the cargo continues to lysosomes for degradation. Hence, anyone of these lysosomal sorting receptors can in theory be used as shuttles for targeted protein degradation.

LDLR, LRP2, and CI-M6PR are all low affinity lysosome sorting receptors for highly abundant ligands (30-32). In contrast, sortilin is a high affinity receptor for low abundance ligands, and specifically the secreted protein progranulin. Sortilin belongs to the Vps10p domain receptor family, and family members show structural and functional resemblance to both LDLR-related receptors and CI-M6PR (33, 34). In fact, a CI-M6PR/sortilin chimera encompassing the sortilin cytoplasmic domain rescues the transport of newly synthesized lysosomal enzymes in CI-M6PR KO cells, suggesting that the two receptors employ overlapping lysosomal sorting routes (35). The sortilin Vps10p domain has been reported to bind several different ligands in vitro including inflammatory cytokines but marked impact on extracellular levels in vivo is only observed for progranulin. The association between sortilin and progranulin is supported by human genetics and Carrasquillo et al. performed a genome-wide screen to identify single nucleotide polymorphisms (SNPs) impacting progranulin plasma level and identified two SNPs located in SORT1 encoding sortilin as the only gene to reach genome-wide significance (12). Accordingly, progranulin levels in plasma and brain are markedly increased in sortilin-deficient mice (11). This effect appears to be highly specific for progranulin as sortilin binds both IL6 and progranulin with single digit nM affinity constants as measured using SPR, but plasma levels of IL6 are slightly reduced in sortilin KO mice (36). The progranulin/sortilin protein-protein interaction is remarkably simple with an absolute requirement for the C-terminal RQLL, apparently constituting a tetrapeptide lysosome targeting tag. Accordingly, we show that extending the C-terminal with RQLL of potentially any protein induces sortilin-mediated endocytosis and degradation.

We further demonstrate that reshaping the progranulin/sortilin interaction can be used as a design framework for plug-and-play generation of targeted degraders of extracellular proteins, denoted SORtilin-based lysosome TArgeting Chimeras (SORTACs). The degrader activity can be genetically encoded as exemplified by converting an IgG-binding nanobody to an IgG degrader by introducing the progranulin C-terminal RQLL, or introduced by chemical conjugation as shown for the therapeutic mAbs alirocumab, adalimumab, and secukinumab, converting them to potent degraders of their targets PCSK9, TNFa, and IL-17A, respectively. Thus, we speculate that a sortilin binding tetrapeptide can be incorporated into most peptide or protein therapeutic drug discovery platform, eliminating the need to identify hits with inhibitory activity on their own.

Finally, we show that small molecules occupying the RQLL binding site in sortilin linked to small molecule target binders generate heterobifunctional degraders with similar size and physicochemical properties as PROTACs (2) but driving targeted degradation of extracellular proteins. Like PROTACs, SORTACs also display the characteristic hook effect and potentially also catalytic activity, evident by the sustained SORTAC-mediated cellular uptake and degradation of the target over time. Hence, the SORTAC technology has the potential to provide a breakthrough in small molecule drug discovery against extracellular targets, eliminating the need for inhibitory activity of the target binder and converting the pharmacology from occupancy-driven to event-driven. As such, conjugation to a sortilin binder can immediately provide functionality to inactive binders discovered in e.g. fragment-based screens or DNA-encoded library screens.

The high expression of sortilin in the peripheral and central nervous system (37) renders SORTACs attractive for targeting neuronal disease proteins such as protein aggregates, neuronal autoantibodies, or inflammatory cytokines. Indeed, we observe that a TNFa targeting SORTAC potently drives the cytokine for degradation in Schwann cells and macrophages. Schwann cells constitute the major glia cell population in the peripheral nervous system and are responsible for myelination of nerve fibers. This is interesting as the interplay between TNFa, Schwann cells and macrophages is central in regeneration following peripheral nerve injury as well as in the development of neuropathic pain (38-40). Furthermore, macrophages play a key role in general tissue inflammation and damage (41), including neuroinflammation (42, 43), and illustrating a wide therapeutic potential of SORTACs.

In conclusion, we here describe the SORTAC platform for the rapid generation of heterobifunctional small molecule degraders against any extracellular protein, expanding the reach of small molecule TPD from the cytosol to the extracellular space.

## Methods

### Chemical synthesis and analysis

Structures, synthesis, and analysis are provided in the Supplementary Methods.

### Cell cultures

All cells were cultured in T25 or T75 flasks (Thermo Fisher) at 37 °C in humidified atmosphere with 5% CO_2_. HEK293 WT (DSMZ, ACC 305), HepG2 (ATCC, HB-8065) and RN22 (Merck, 93011414) were cultured in DMEM (Lonza) supplemented with 10% heat in-activated fetal bovine serum (FBS) (Sigma-Aldrich), 1% penicillin-streptomycin (P/S) (Sigma-Aldrich), 1% GlutaMAX (Gibco). HEK293/sortilin (44) were cultured in corresponding medium supplemented with 100 μg/mL Zeocin (Invitrogen). K562 (ATCC, CCL-243) were cultured in RPMI (VWR) + 10% FBS + 1% P/S + 1% GlutaMAX. SH-SY5Y (ATCC, CRL-2266) were cultured in DMEM:F12 (Gibco) + 10% FBS + 1% P/S + 1% GlutaMAX. Sub-cultivation of adherent cells was performed using 2.5% trypsin (Gibco) + 0.1 mM EDTA (Thermo Fisher). Cell lines were authenticated by the supplier. Cell lines were verified to be mycoplasma free using MycoplasmaCheck (Eurofins Genomics) or by the supplier. Mouse CD1 Brain Cortex Neurons (Lonza, cat.nr.: M-CX-400) were thawed, seeded and cultured according to manufacturer’s protocol.

Primary Schwann cell cultures were prepared from neonatal Sprague Dawley wild type (Janvier labs) and Sort1^-/-^ rats (45). In brief, sciatic nerves were dissected from P1-P3 pups and placed in ice-cold Leibovitz’s L-15 medium (Gibco). Following dissection, nerves were digested with 0.25% trypsin (Gibco) and 0.1% collagenase (Sigma-Aldrich) for 30 min at 37°C and dissociated by trituration in DMEM (Sigma-Aldrich) containing 10% FBS (Thermo Fisher). Cells were subsequently plated on poly-L lysine-coated culture dishes in DMEM with 10% FBS and Primocin (InvivoGen). Fibroblasts were eliminated by incubation with the anti-metabolic agent Cytosine b-D-arabinofuranoside (Ara-C; 10 µM final concentration, Sigma-Aldrich) for 48 hours (h). Purified primary Schwann cells where then expanded in growth medium (DMEM supplemented with 10% FBS, 1% P/S recombinant human neuregulin1-b1/HRG1-b1 EGF domain (10 ng/ml, R&D Systems) and forskolin (2.5 µM, Sigma-Aldrich)) at 37°C, 5% CO_2_. Peripheral blood mononuclear cells (PBMCs) were isolated from healthy donors through Ficoll-Paque PLUS (Cytiva 17-1440-03) using SepMate density gradient centrifugation tubes (StemCell Technologies, 85450). For macrophage differentiation, monocytes were isolated from PBMCs by negative selection (STEMCELL technologies; 19059) and cultured for 5 days at 37 °C in 96-well plates in the presence of M-CSF (Biolegend, 574804) in RPMI 1640 media containing 10% FBS and 1% P/S.

### Mammalian protein expression

#### Plasmids

Sortilin extracellular domain (ECD) (M1-S756, Acc: CAA66904.2) and PCSK9 (Acc. CAC38896.1) both C-terminally fused to 6His (Sortilin-6His, PCSK9-6His), PCSK9 C-terminally extended with (GGGGS)3GGRQLL) (PCSK9-RQLL), IL-17A (Acc. NP_002181.1) N-terminally tagged with NanoLuc® (Promega) (NanoLuc®-IL-17A) and progranulin (NP_002078.1) N-terminally tagged with the HiBiT peptide (Promega) (HiBiT-PGRN) were all expressed using the pCpGfree-vitroNmcs vector (InvivoGen).

#### Expression

Sortilin-6His, NanoLuc®-IL-17A and HiBiT-PGRN were produced using the ExpiCHO expression system (46). PCSK9-6His and PCSK9-RQLL stably transfected in CHO-K1 cells were adapted to growth in suspension in Hybridoma-SFM medium (GIBCO) and then expanded in Celline CL100 Bioreactor flasks (Integra).

#### Purification

The pH of Sortilin-6His medium was adjusted to 7.4 by adding Tris-HCl to 50 mM. Imidazole was added to 10 mM before loaded onto a 5 mL HisTrap Excel column (Cytiva) equilibrated in 50 mM Tris-HCl pH 7.4, 500 mM NaCl, 10 mM imidazole. The loaded column was washed to baseline and protein of interest was eluted with 400 mM imidazole. Protein was dialysed (MWCO: 6-8 kDa) against PBS overnight (ON) at 4 °C followed by concentration in a centrifugal concentrator (MWCO: 30 kDa). The concentrated sample was subjected to size-exclusion chromatography on a Superdex200 Increase column (Cytiva) equilibrated in PBS.

The pH of PCSK9-6His medium was adjusted to 7.4 by adding Tris-HCl to 50 mM and loaded onto a 5 mL HisTrap Excel column (Cytiva) equilibrated in 50 mM Tris-HCl pH 7.4, 500 mM NaCl, 20 mM imidazole, 1 mM CaCl2. The loaded column was washed to baseline and protein of interest was eluted with 400 mM imidazole. Protein was dialysed (MWCO: 14 kDa) against 25 mM Tris-HCl pH 7.4, 150 mM NaCl, 5 % glycerol ON at 4 °C followed by concentration in a centrifugal concentrator (MWCO: 30 kDa). The concentrated sample was subjected to size-exclusion chromatography on a Superdex200 Increase column (Cytiva) equilibrated in 25 mM Tris-HCl pH 7.4, 150 mM NaCl.

PCSK9-RQLL medium was added Tris-HCl pH 7.4 to a final concentration of 25 mM before loaded onto a 5 mL HiTrap QFF column (Cytiva) equilibrated in 50 mM Tris-HCl pH 7.4. The loaded column was washed to baseline and protein of interest was eluted with 25 mM Tris-HCl pH 7.4, 1 M NaCl. Purified protein was dialysed (MWCO: 14 kDa) against 25 mM Tris-HCl pH 7.4, 5% glycerol ON at 4 °C followed by concentration in a centrifugal concentrator (MWCO: 30 kDa). The concentrated sample was loaded onto a 5 mL HiTrap Heparin HP column (Cytiva) equilibrated in 25 mM Tris-HCl pH 7.4. Protein was incubated for 30 min on-column before being washed to baseline and eluted with a linear gradient from 0-650 mM NaCl and a final 1 M NaCl elution step. Fractions containing protein of interest were pooled and concentrated in a centrifugal concentrator (MWCO: 30 kDa) and subjected to size-exclusion chromatography on a Superdex200 Increase column (Cytiva) equilibrated in 25 mM Tris-HCl pH 7.4, 150 mM NaCl.

HiBiT-PGRN medium was diluted 1:5 in 50 mM Tris-HCl pH 7.4 before loaded onto a 5 mL HiTrap QFF column (Cytiva) equilibrated in 50 mM Tris-HCl pH 7.4. FT was collected, concentrated in a centrifugal concentrator (MWCO: 30 kDa) and subjected to size-exclusion chromatography on a Superdex200 Increase column (Cytiva) equilibrated in PBS.

Purified proteins were concentrated, flash-frozen and stored at -80 °C.

NanoLuc®-IL-17A media was clarified by centrifugation and supernatant was used as conditioned media.

### *E. coli* protein expression

#### Plasmids

Plasmids expressing TNFa (V76-L233 Acc. NP_000585.2), Enhanced GFP (Acc. U55762), and anti-kappa light chain (LC) specific V_H_H nanobody (Nb) (PMID: 35551129) encoding N-terminal 6His tag and TEV protease recognition sites and C-terminal (GGGGS)3GG linkers with and without the C-terminal RQLL motif were cloned into the pET11d vector (NOVAGEN). In addition, TNFa (V76-L233) was cloned in the same vector without a C-terminal tag.

#### Expression

Plasmids were transformed into the *E. coli* BL21(DE3) or Shuffle-T7-Express-Lys-Y (NEB) and plated on LB plates containing 100 µg/mL ampicillin following incubation at 37 °C for 18 h. Overnight cultures were grown in 100 mL LB medium (Luria–Bertani) at 30 °C and 100 µg /mL ampicillin at 175 RPM. Cell pellets from overnight cultures were resuspended in 1 L of 2xYT medium (Tryptone 16 g/L, Yeast Extract 10 g/L, NaCl 5 g/L, 100 µg/mL ampicillin) and incubated at 30 °C at 200 RPM. At an OD600 of 0.75, Isopropyl ß-D-1-thiogalactopyranoside (IPTG) was added to a final concentration 0.4 mM. Cultures were then incubated at 16 °C or 19 °C at 200 RPM for 18 h. Cell pellets were harvested at 6000 RPM for 15 min and kept at -20 °C.

#### Purification

Pellets of cells expressing TNFa+/-RQLL were resuspended in 20 mM Tris-HCl pH 8.0, 500 mM NaCl, 20 mM imidazole, 1 mM PMSF and lysed by sonication. The lysate was clarified by centrifugation and supernatant was loaded onto a 5 mL HisTrap FF Crude column (Cytiva) equilibrated in 20 mM HEPES pH 7.5, 500 mM NaCl, 20 mM imidazole. The loaded column was washed to baseline and elution was performed with step gradient from 20-300 mM imidazole. Protein was dialysed (MWCO: 6-8 kDa) against 20 HEPES pH 7.5, 150 mM NaCl ON at 4 °C followed by concentration in a centrifugal concentrator (MWCO: 5 kDa). The concentrated sample was subjected to size-exclusion chromatography on a Superdex75 10/300 Increase column (Cytiva) equilibrated in 25 mM HEPES pH 7.5, 150 mM NaCl. Enhanced GFP+/-RQLL pellets were lysed by sonication and centrifuged 40 min at 12000 rpm before filtering the supernatant and applying it to a pre-equilibrated 5 ml HisTrap HP column (Cytiva). The column was washed with 4 column volumes (CV) of 50 mM Tris pH 7.5, 300 mM NaCl and 20 mM imidazole and proteins were eluted with 3 CV 50 mM Tris pH 7.5, 300 mM NaCl and 300 mM imidazole. Eluted proteins were treated with 6xHIS-TEV protease and dialyzed against PBS ON at 4 °C. The dialyzed sample was applied to a 5 ml HisTrap HP column and untagged proteins were subsequently collected in flow-through and wash fractions. Untagged proteins were finally applied to a SD200 increase 10/300 (Cytiva) gel-filtration column in PBS and eluted fractions were analyzed by SDS-PAGE. Nb+/-RQLL pellets were resuspended in 20 mM Tris-HCl pH 8.0, 500 mM NaCl, 20 mM imidazole, 1 mM PMSF and lysed by sonication. The lysate was clarified by centrifugation and supernatant was loaded onto a 5 mL HisTrap FF Crude column (Cytiva) equilibrated in 20 mM Tris-HCl pH 8.0, 500 mM NaCl, 20 mM imidazole. The loaded column was washed to baseline and elution was performed with a linear gradient from 20-300 mM imidazole. Protein was dialysed (MWCO: 6-8 kDa) against 20 mM Sodium acetate pH 5.5, 50 mM NaCl ON at 4 °C followed by concentration in a centrifugal concentrator (MWCO: 5 kDa). The concentrated sample was subjected to size-exclusion chromatography on a Superdex75 Increase column (Cytiva) equilibrated in 25 mM Tris-HCl pH 7.4, 150 mM NaCl. The TNFa-6His pellet was solubilized in 20 mM Tris-HCl pH 8.2, 500 mM NaCl, 10 mM imidazole, 1 mM PMSF (resuspension buffer) and sonicated for 3 x 5 min at 40% amplitude. Lysed cells were centrifuged at 18514 x g for 20 min to remove cell debris and supernatant was filtered through an 0.8 µm filter and loaded onto a 5 mL HisTrap HP column (Cytiva) equilibrated in resuspension buffer. The loaded column was washed to baseline and protein of interest was eluted with 400 mM imidazole. Protein was dialyzed (MWCO: 6-8 kDa) against PBS ON at 4 °C before being concentrated in a centrifugal concentrator (MWCO: 10 kDa) and subjected to size-exclusion chromatography on a Superdex75 Increase column (Cytiva) equilibrated in PBS. Pure protein samples were concentrated, flash-frozen and stored at -80 °C.

### Peptides

Peptidic degraders and NHS-peptides for conjugation of antibodies were synthesized by WuXi AppTec.

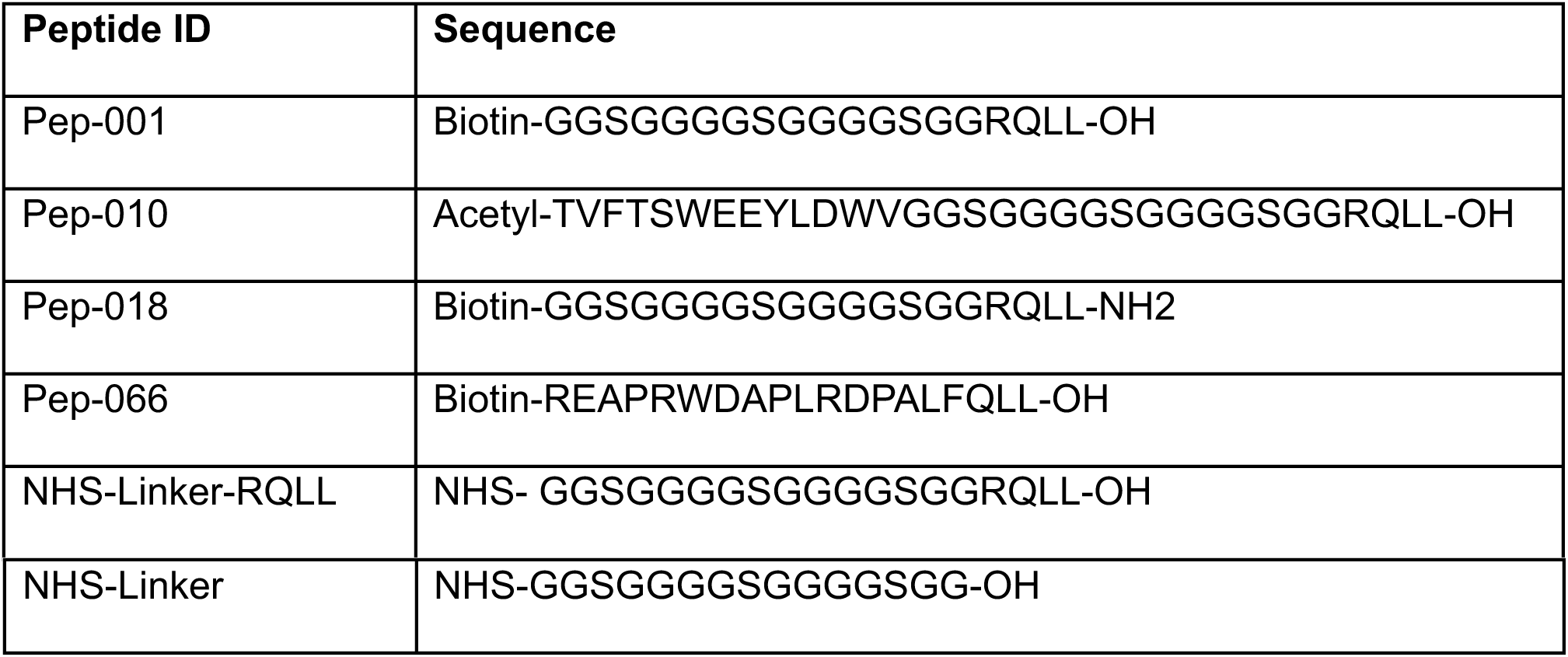

Pep-066 has the G/S linker replaced by an extended progranulin C-terminal sequence stretch and an optimized sortilin binding tetrapeptide (FQLL) (data not shown).

### Chemical conjugation of proteins

Proteins (TNFa-6His, TNFa-6His-RQLL, IgG/kappa LC (alirocumab), PCSK9-6His) in PBS were mixed in a 1:8 molar ratio with Cyanine5 (Cy5) NHS ester (Lumiprobe) or pHrodo™ Red (pHrodo) succinimidyl ester (Invitrogen) and incubated 1 h at room temperature (RT). Labelled proteins were purified from unreacted Cy5 or pHrodo esters using Desalting Gravity Flow Column (BioRad) with PBS as buffer.

Antibodies (alirocumab/adalimumab/secukinumab) in PBS (4 mg/ml) were mixed in a 1:25 molar ratio with NHS-Linker-RQLL or NHS-Linker peptide and incubated ON at RT. Labelled antibodies were purified from unreacted NHS-peptide using a desalting column (Thermo Fisher) with PBS as buffer. The linkage reaction resulted in a shift of approximately 8-12 kDa in molecular weight of the heavy chain and light chain fragments of the antibody as visualized by reducing SDS-PAGE and indicating that each antibody was linked at 2-10 positions (Extended Data Fig 4e).

### Microscale Thermophoresis

Binding to sortilin ECD was assessed by Microscale Thermophoresis (MST). In these experiments a 24- or 12-point-titration series of ligand solution was prepared in TTP LVDS 384-well plates (SPT Labtech) in 1.2 µL total volume using a Mosquito pipetting robot (SPT Labtech) in a buffer composed of 16.7% DMSO (v/v) and 0.05% Tween20 (v/v). The plate was spun before addition of 8.8 µL solution containing 114 nM sortilin ECD and 28.4 nM RED-tris-NTA (Nanotemper) in 57 mM Bis-Tris Propane pH 9, 57 mM NaCl, 0.05% Tween20 (v/v) to each well. Reaction was incubated 1 h at 19 °C. MST was measured on a Monolith NT. Automated (Nanotemper) in standard-treated capillaries at 5% LED-power and high MST-power. Dose-response was extracted from the raw thermographs at 20 sec hot-time.

Additionally, MST was performed to assess affinity of sortilin to GFP-RQLL and GFP. In these experiments a 12 point-titration series of sortilin was prepared in PCR tubes in 5 µL total volume using 25 mM HEPES pH 7.5, 150 mM NaCl og 0.05 % Tween20 as buffer. 5 µL protein solution containing 40 nM GFP-RQLL or 10 nM GFP in 25 mM HEPES pH 7.5, 150 mM NaCl and 0.05 % Tween20 was added to each well. Reaction was incubated 1 h at 19 °C. MST was measured on a Monolith NT. 115 (Nanotemper) in standard-treated capillaries at 20% LED-power and medium MST-power. Dose-response was extracted from the raw thermographs at 2.5 sec hot-time.

Dissociation constants were estimated by sigmoidal curve fitting in MO. Affinity analysis program (Nanotemper).

### Size exclusion chromatography for analysis of complex formation

Size exclusion chromatography was used to confirm target binding of nanobody and mAb degraders. Kappa LC specific Nb containing the RQLL motif (Nb-RQLL) or linker without RQLL motif (Nb-Linker) was mixed with IgG/kappa LC (alirocumab) in a 2.2:1 molar ratio in a final volume of 250 µL and incubated for 1 h at RT. Samples were centrifuged for 15 min at 13400 rpm at 4 °C to remove potential aggregates and applied to a Superdex75 Increase 10/300 connected to a Äkta Pure HPLC system (Cytiva). A control sample using Nb-RQLL or Nb-Linker alone was analysed for comparison.

Alirocumab containing the RQLL motif (Ali-RQLL) or linker without RQLL motif (Ali-Linker) was mixed with PCSK9 in a 1:2.5 molar ratio in a final volume of 200 µL and incubated ON at 4 °C. The complex sample was centrifuged as described above and applied to a Superdex200 Increase 10/300 (Cytiva). A control sample using PCSK9 was analysed for comparison.

For all experiments, 0.5 mL fractions were collected and peak fractions containing the complexes were analysed by SDS-PAGE/Coomassie staining.

### Cellular uptake assay

For assessment of cellular uptake, cell lines (HEK293/sortilin, HEK293, HepG2, RN22, K562 and SH-5YSY) and primary cells (WT and sortilin KO rat Schwann cells) were seeded in poly-L-Lysine coated 96-well plates (Costar) and incubated ON, before replacement of culture medium with fresh medium containing the target of interest (NeutrAvidin-650 (NA650) (Invitrogen), Cy5-IgG (alirocumab), Dinitrophenyl-KLH polyclonal rabbit antibody conjugated to Alexa Fluor™ 488 (anti-DNP-AF488) (Invitrogen), Cy5-TNFa, Cy5-TNFa-RQLL, pHrodo-TNFa, Cy5-PCSK9, Cy5-PCSK9-RQLL, NanoLuc®-IL-17A) and SORTACs as indicated. Leupeptin (Merck), monofunctional sortilin binders (AF38469, MSM, MSM2, pep-010) or target binders were added as indicated. Following incubation (2-24 h), cells were washed three times in PBS and the fluorescence signal was quantified using a plate reader (BMG Labtech, Clariostar). For NanoLuc®-IL-17A, cells were lysed in 50 µL Passive Lysis Buffer (Promega) followed by the addition of 50 µL Nano-Glo® Luciferase Assay System substrate (Promega) (47). Luciferase activity was evaluated by luminescence measurements at 470 nm using a plate reader (Clariostar, BMG Labtech).

### Confocal microscopy

HEK293/sortilin cells were seeded on poly-L-Lysine coated coverslips (25K/coverslip) and incubated ON before replacement of medium with culture medium with 500 nM NA650 (Invitrogen) and 2 µM pep-001 with or without 10 µM AF38469 (MedChemExpress). Following 4 h incubation, cells were washed in PBS and fixed for 15 min in 4% PFA (Sigma-Aldrich), permeabilized for 15 min with 0.1% Triton X-100 (Sigma-Aldrich) in tris-buffered saline (TBS), blocked for 20 min in 2% BSA (Sigma-Aldrich) in TBS and washed in TBS before incubation with anti-LAMP1 antibody at 4 °C ON. Cells were washed in TBS and incubated with goat-anti-mouse AF488 for 2 h at RT before they were washed again and incubated in Hoechst diluted 1:10000 (Sigma-Aldrich) for 10 min. Nb-RQLL activity was evaluated in a similar manner; cells were incubated with 500 nM Cy5-IgG and 250 nM Nb-RQLL or Nb control. Following 3 h incubation, cells were washed and fixed before Hoechst staining. Mouse CD1 cortex neurons were seeded 80K cells on poly-D-Lysine and laminin coated coverslips in culture medium. After 13 days in culture, medium was replaced with culture medium with 500 nM NA650 (Invitrogen) and 560 nM BSM4 or biotin. Following 3 h incubation, cells were washed and fixed before Hoechst staining. All coverslips were mounted using fluorescence mounting medium (Agilent Dako) before evaluation of fluorescent signal using a laser scanning confocal microscope (LSM-800). In all experiments 3 coverslips of each condition were inspected and a minimum of five representative images were processed using Zeiss ZEN 3.5 Blue edition software and ImageJ. Monocyte-derived macrophages were pre-incubated in the absence or presence of leupeptin (Sigma-Aldrich) prior to incubation in the absence (vehicle) or presence of either: BSM14, BSM16 or MSM3 (at the indicated concentrations) in the presence of Cy5-TNFa for 4 h. After the culture period, cells were labelled using CellTrace Calcein Green, AM (Thermo Fisher) and Hoechst followed by image acquisition by confocal microscopy (Yokogawa CQ1).

### Protein degradation analysis by Western blotting

HEK293/sortilin cells were seeded in poly-L-Lysine coated plates (250K/24-well) and incubated ON, before incubation in medium containing target (Cy5-IgG, anti-DNP-AF488 (Invitrogen), TNFa-6His), SORTACs, sortilin and target monobinders and leupeptin (Merck) as indicated. Following incubation, cells were washed in PBS and lysed 30 min on ice in STE buffer (Sigma-Aldrich) with 1% Nonidet™ P 40 Substitute (Sigma-Aldrich) and cOmplete (Roche). To study the time course of target degradation, target and degrader containing medium was removed after initial uptake-incubation, and cells were washed in PBS before addition of fresh medium and further incubation before harvest of cells at indicated time-points. Samples were spun and prepared with LDS sample buffer (Thermo Fisher) and 17 mM DTT (Sigma-Aldrich) and boiled for 5 min before subjected to SDS-PAGE using a 4-12 % Bis-Tris gel (Thermo Fisher). Proteins were blotted onto a nitrocellulose membrane (Thermo Fisher). Fluorescently labelled targets (NA650, Cy5-IgG/kappa LC, anti-DNP-AF488), were detected using an iBright 1500 (Thermo Fisher). Next, membrane was blocked for 1 h in blocking buffer (TBS + 1% Tween20 (TBS-T) (Sigma-Aldrich) and 5 % milk powder), and incubated with primary antibody in blocking buffer for 2 h at RT or ON at 4 °C. The membrane was washed 3 times in wash buffer (TBS-T + 0.5 % milk powder) and incubated with HRP conjugated secondary Ab for 1 h at RT before it was washed 3 times in wash buffer. Blot was developed and analyzed using ECL reagents (Cytiva) and iBright 1500 (Thermo Fisher). Quantification of band intensity was done using iBright analysis software (Thermo Fisher). Selected membranes were stripped in Restore stripping buffer (Thermo Fisher) for 15 min at 37 °C and reprobed following the procedure for blocking and antibody incubation described above.

### Extracellular depletion assay

HEK293/sortilin cells were seeded in poly-L-Lysine coated plates (40K/96-well) and incubated ON before replacement of culture medium with medium containing target (NA650 (Invitrogen), Cy5-IgG, aDNP IgG (Thermo Fisher) or TNFa) and degrader molecules as indicated. For fluorescently labelled targets, FluoroBright DMEM (Gibco) supplemented with 10% FBS (Sigma-Aldrich) and 1% Glutamax Supplement (Gibco) were used. After incubation for up to 72 h, remaining target in cell culture supernatant was evaluated by in blot analysis of fluorescence intensity (iBright 1500, Thermo Fisher) following SDS-PAGE and transfer of proteins to a nitrocellulose membrane (Cy5-hIgG) or by fluorescence intensity measurements using a plate reader (Clariostar, BMG Labtech) (NA650). For TNFa and aDNP IgG, the remaining target in medium was assessed by anti-human TNFa ELISA (Biolegend, 430201) or anti rat IgG ELISA (Invitrogen, 88-50490-22) according to manufacturer’s protocol.

### Antibodies

Antibodies used in the study are listed in table. Detailed information on antibodies used in WB and IF can be found in the pAbmAbs research antibody review database www.pAbmAbs.com deposited by Jonas Lende, Ditte Køster and Marianne Lundsgaard Kristensen.

| Ag | Clonality | Species | labeling | Application | Cat# | Source |
| --- | --- | --- | --- | --- | --- | --- |
| β actin | monoclonal | mouse |  | WB<br>(1:5000) | A5441 | Sigma-Aldrich |
| sortilin | Monoclonal | mouse |  | WB<br>(1:2000) | 612100 | BD Bioscience |
| Human<br>IgG | monoclonal | rabbit |  | WB<br>(1:2000) | EPR12700<br>ab181236 | Abcam |
| TNFA | monoclonal | mouse |  | WB<br>(1:2000) | MA5-23720 | Invitrogen |
| Rabbit<br>IgG | polyclonal | goat | HRP | WB<br>(1:5000) | ab7090 | Abcam |
| Mouse<br>IgG | polyclonal | goat | HRP | WB<br>(1:5000) | ab6789 | Abcam |
| LAMP1 | monoclonal | mouse |  | IF (1:200) | H4A3<br>Ab25630 | Abcam |
| Mouse<br>IgG | polyclonal | donkey | AF488 | IF (1:500) | A-21202 | Invitrogen |
| 6-HIS | monoclonal |  | Tb<br>Cryptate | TR-FRET (2<br>nM) | 61HI2TLA | Revvity |
| DNP | polyclonal | rabbit | AF488 | In vitro<br>assay | A-11097 | Thermo Fisher |
| DNP | monoclonal | rat |  | In vitro<br>assay | 04-8300 | Thermo Fisher |
| PCSK9 | monoclonal | humanized |  | In vitro<br>assay | alirocumab | Sanofi/Regeneron |
| TNFα | monoclonal | humanized | | In vitro<br>assay | adalimumab | Abbvie |
| IL-17A | monoclonal | humanized |  | In vitro<br>assay | secukinumab | Novartis |

### Time-resolved fluorescence resonance energy transfer (TR-FRET)

Ternary complex formation between the target of interest, SORTAC and sortilin ECD was quantified using TR-FRET. SORTACs, biotin or antibodies were diluted in a 384-well TTP LVDS plate (SPT Labtech) to a final TR-FRET buffer composition of 50 mM Bis-Tris Propane pH 8, 50 mM NaCl and 0.05% Tween20, with or without 10% DMSO using a Mosquito LV pipetting robot (SPT Labtech). In a black 384-well plate (Corning) the dilutions were mixed with following components in TR-FRET buffer to final concentrations of donor: 2 nM Tb cryptate-labelled anti-6His antibody (mAb Anti-6His Tb, Revvity), acceptor: 50 nM d2-labelled Streptavidin (Streptavidin-d2, Revvity), donor protein: 40 nM, 100 nM or 200 nM His-tagged sortilin ECD and for protein targets a biotin-labelled acceptor protein (40 nM PCSK9-Biotin or 150 nM TNFa-Biotin). The reactions were incubated for 2.5 h at RT. The HTRF signal ratio at 665/620 nm was determined using a plate reader (ClarioStar, BMG Labtech).

### Data Analysis

MARS data analysis software (BMG Labtech) was used for pre-processing of raw plate reader data and absolute quantification of ELISA data from standard using 4-parameter nonlinear regression. Further data-handling was performed in Prism (Graph-pad).

Cell uptake and extracellular depletion data are displayed as background corrected raw values or increase in raw values (FI or Luminescence) above baseline or as normalized values (%FI). Normalized values are calculated as: (FI - background)/(baseline - background). Background is defined by signal from wells without addition of target. Baseline is defined by signal from wells with target, but without SORTAC. For assays measuring inhibition of the cellular uptake, baseline is defined by wells with target and SORTAC, but without inhibitor.

Gaussian non-linear regression of data curves dependent on ternary complex formation were used to calculate amplitude, ternary complex-forming EC_50_ (TF_50_), ternary complex– inhibitory EC_50_ (TI50) and maximally effective concentration (ECmax) complex forming-max (TFmax) as described in (48). p-values were calculated using unpaired two-tailed t-tests.

All data shown is representative of at least three independent experiments of each at least two biological replicates.

### Structural biology

The small molecule SORTAC BSM15 targeting TNFa was dissolved to 20 mM in 100% DMSO and mixed with purified sortilin ECD (6.6 mg/mL) in a 1:20 v/v ratio, ie. to a final compound concentration of 1 mM and 5% DMSO. The SORTAC/sortilin complex was subsequently mixed with crystallization buffer containing 22% PEG6000, 1.1 mM NaCl, 8% glycerol, 50 mM HEPES pH 8.0 in a 1:1 ratio and co-crystals were obtained using the sitting-drop vapor diffusion method at 19 °C. After 14 days, co-crystals were harvested and flash-frozen in liquid N2. Diffraction images were collected at the BioMAX x-ray beamline at MaxIV Laboratory, Lund, Sweden. Images were processed and scaled using the XDS package (49) and a resolution limit of 3.0 Å (criteria: I/sigma > 1.0). Molecular replacement and initial refinement were performed using DIMPLE within CCP4i2 (50, 51) and coordinates from an isomorphous sortilin ECD x-ray structure (PDB ID: 4PO7) as template. Further refinement, mostly around SORTAC-binding site, was carried out by alternating cycles of manual model building in *Coot* (52) and refinement in REFMAC5 within CCP4i2. Coordinates and geometry restraints for BSM15 were created using AceDRG and manually fitted within the positive electron density in *Coot*. Figures 6 g-h were prepared in PyMOL (The PyMOL Molecular Graphics System, Version 3.0 Schrödinger, LLC).

**Table 1.**
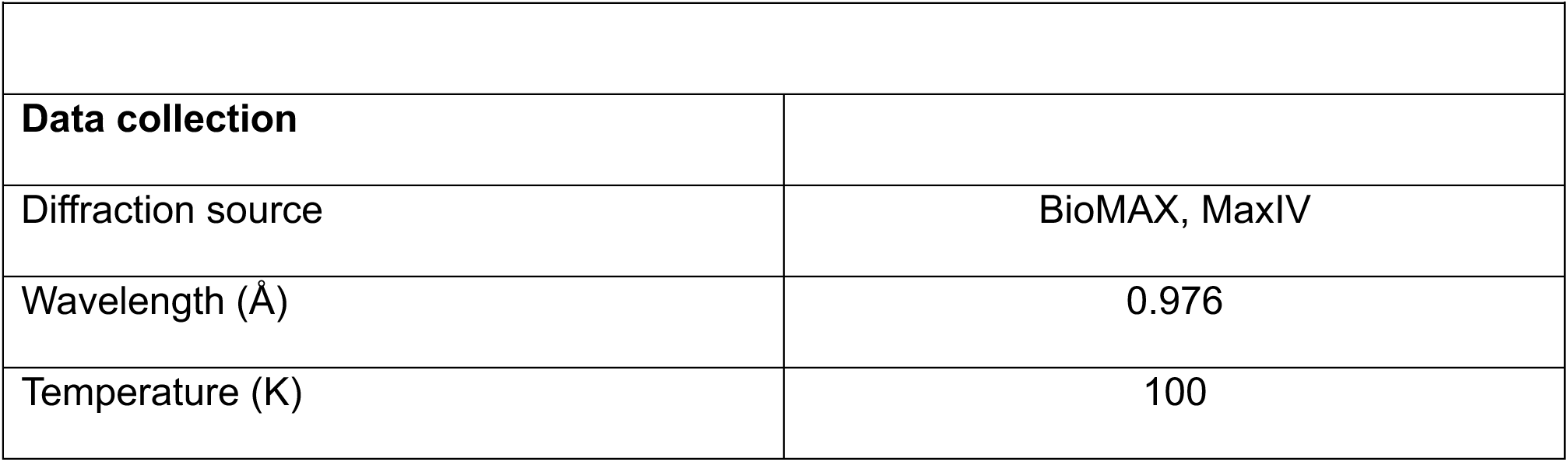
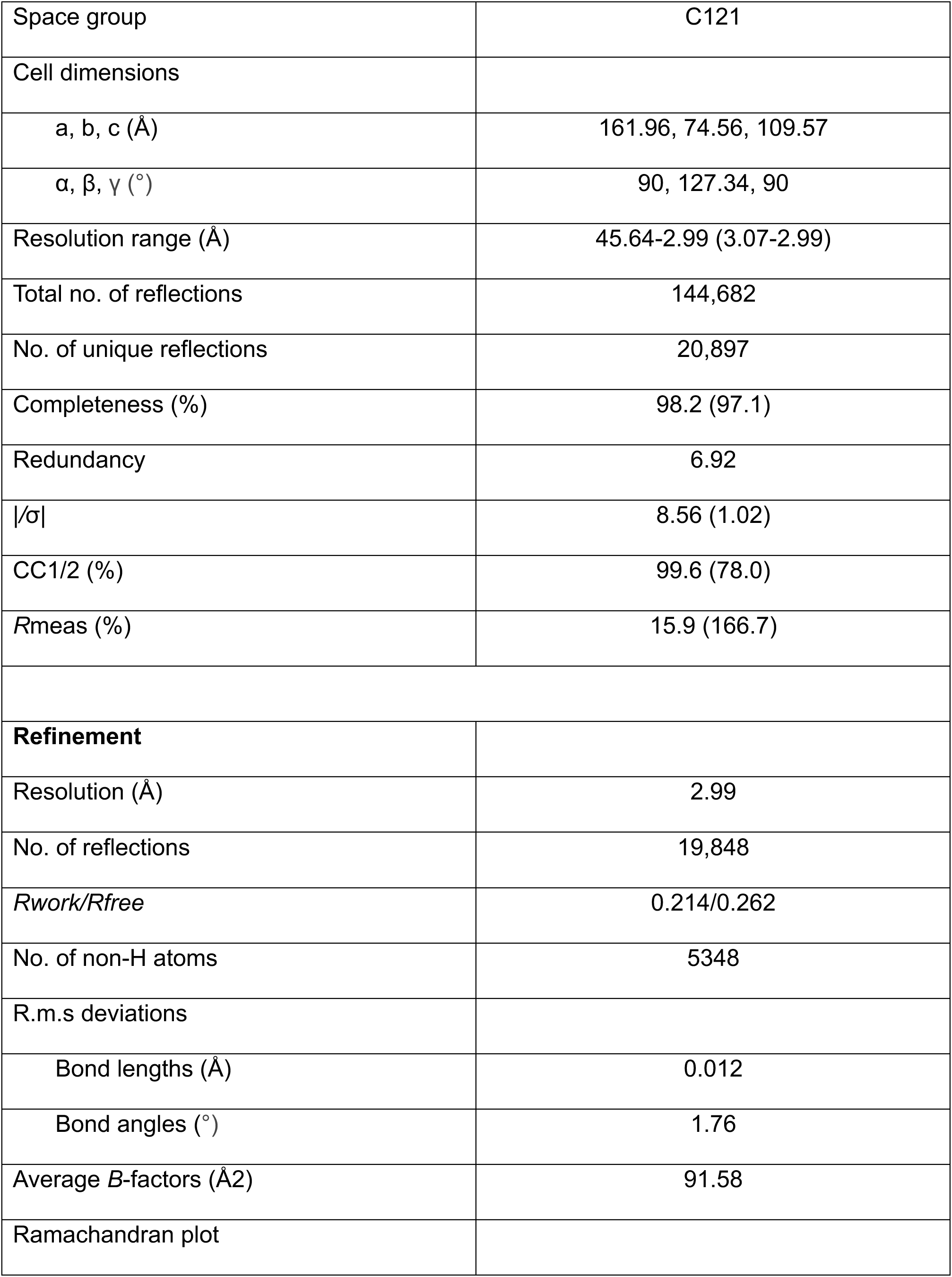
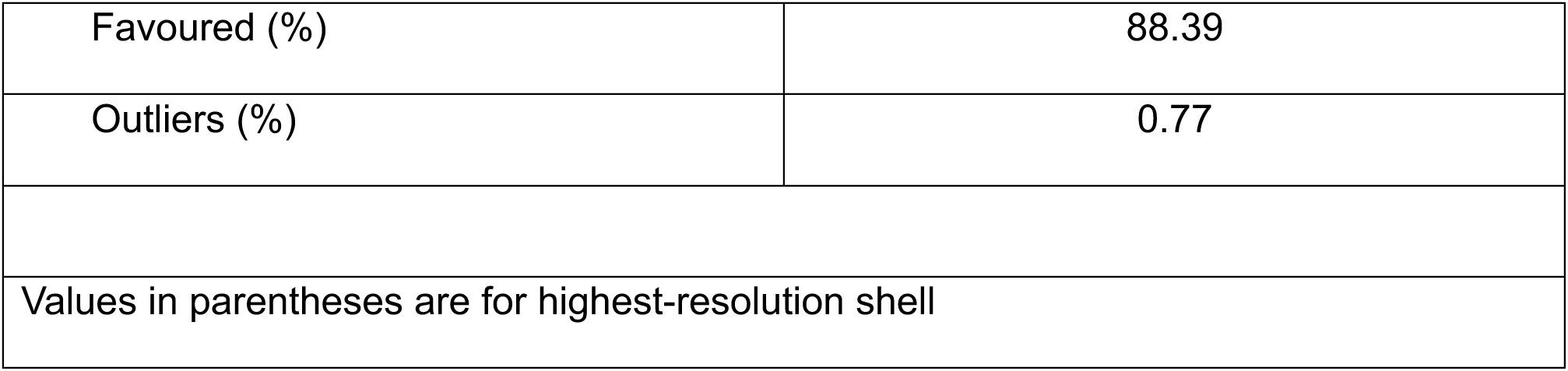
Data collection and refinement statistics.

## Data availability

All data is presented in the manuscript and Supplementary methods section.

## Acknowledgements

The study was funded by Draupnir Bio, the Novo Nordisk Foundation Pioneer Innovator program (NNF20OC0066247), and the Innovation Fund Denmark Innoexplorer and Innobooster programs.

---

**Extended data Fig. 1.**
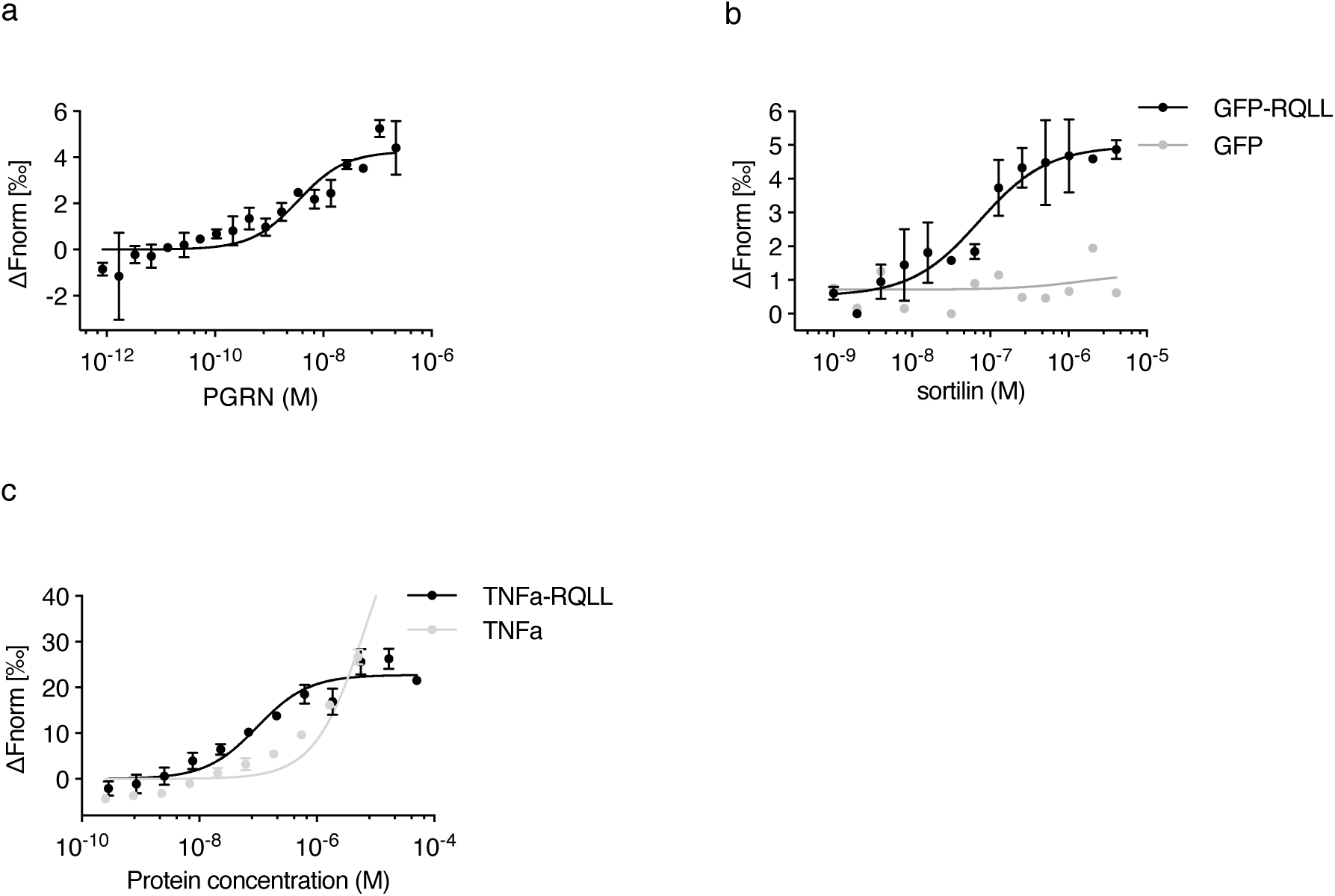
**a-c**, microscale thermophoresis (MST) measurements of sortilin ECD binding to PGRN (**a**), GFP and GFP-RQLL (b), and TNFa and TNFa-RQLL (**c**). **a**, The titration of PGRN (219 nM to 0.8 pM) with a constant concentration of fluorescently labelled sortilin ECD at 100 nM. The change in thermophoretic signal allows calculation of the K_D_ = 3.5 ±1.6 nM (n = 2). **b**, The titration of sortilin ECD from 4.1 µM to 1 nM with a constant concentration of GFP-RQLL or GFP at 20 nM. The change in thermophoretic signal allows calculation of the K_D_ = 74 nM for the interaction with GFP-RQLL, while no binding affinity to GFP could be detected. (n = 2 (GFP-RQLL) and n=1 (GFP)). **c**, The titration of the ligand concentration from 45 µM to 254 pM TNFa or 50.5 µM to 285 pM TNFa-RQLL with a constant concentration of fluorescently labelled sortilin ECD at 100 nM. The change in thermophoretic signal allows calculation of the K_D_ = 86.5 ±48 nM for the interaction with TNFa-RQLL, while the binding affinity to TNFa was defined as >200 µM due to an incomplete curve fit. (n = 4 independent measurements). Error bars represent standard deviation.

**Extended data Fig. 2.**
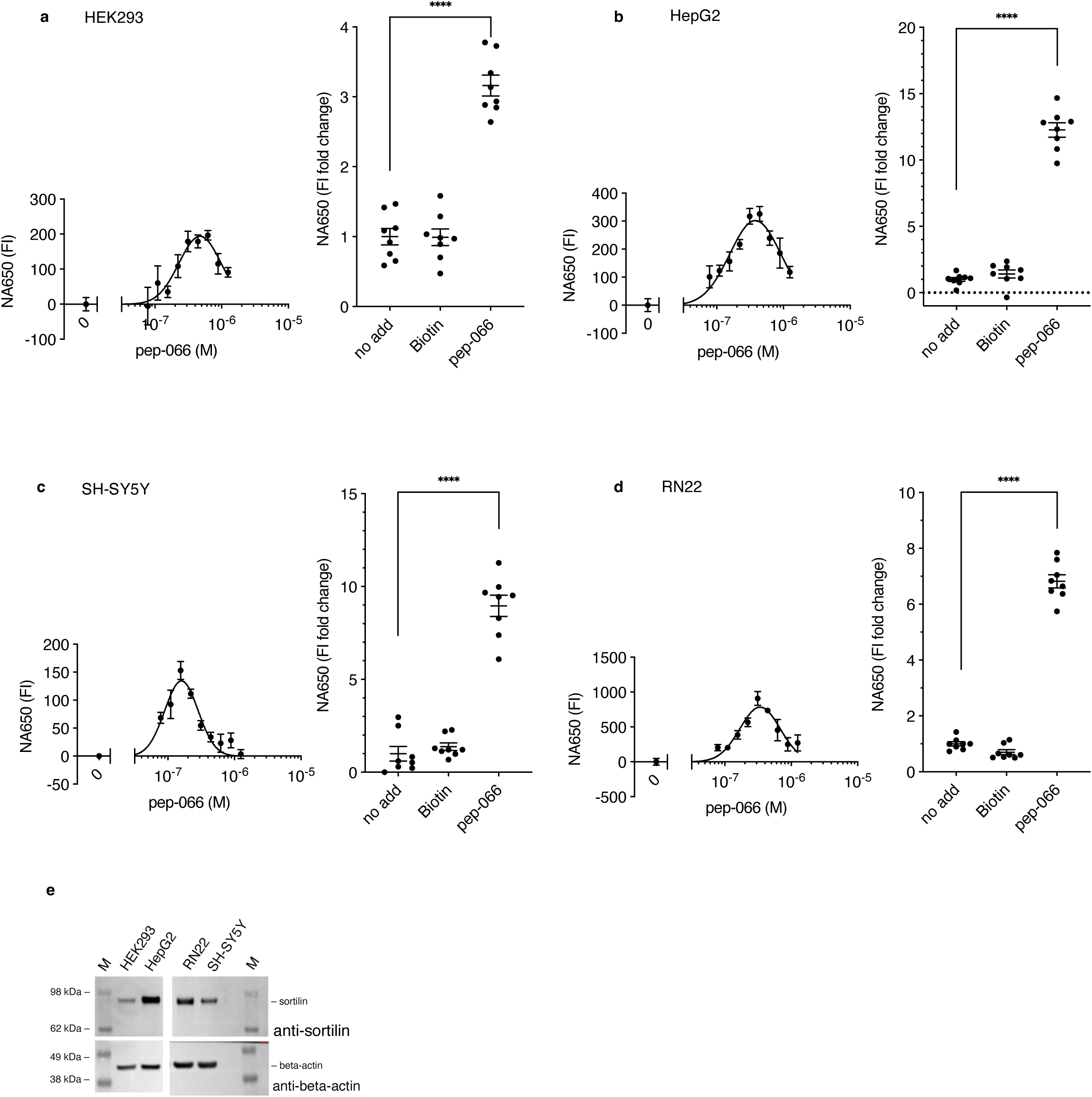
**a-d,** To investigate SORTAC activity in cells with endogenous sortilin expression, cell lines HEK293 (a), HepG2 (b), SH-5YSY (c), and RN22 (d) were seeded in 96-well plates and incubated ON, before replacement of culture medium with fresh medium containing NA650 (100 nM) and a dilution series of pep-066. Following 24 h incubation, cells were washed, and the FI signal was quantified using a plate reader. Data is shown as the increase in FI above baseline (0 nM pep-066). Mean±SEM (n=2). In a separate experiment, cells were incubated (24 h) with NA650 (100 nM) and a single concentration of pep-066 (312 nM). Control cells were incubated with NA650 and target binder biotin (312 nM). Bar graphs show data normalized to mean value of samples without addition of degrader (no add). Mean±SEM (n=8). p-values were determined by unpaired two-tailed t-test: *p<0.05, **p<0.01, ***p<0.001, ****p<0.0001. e, Sortilin expression was confirmed by Western blotting of cell lysates (10 µg total protein loaded on SDS-PAGE gel). Beta-actin blot (lower blot) shown as control. n indicates the number of biological replicates in a given experiment. Each data set shown is representative of at least three independent experiments.

**Extended data Fig 3.**
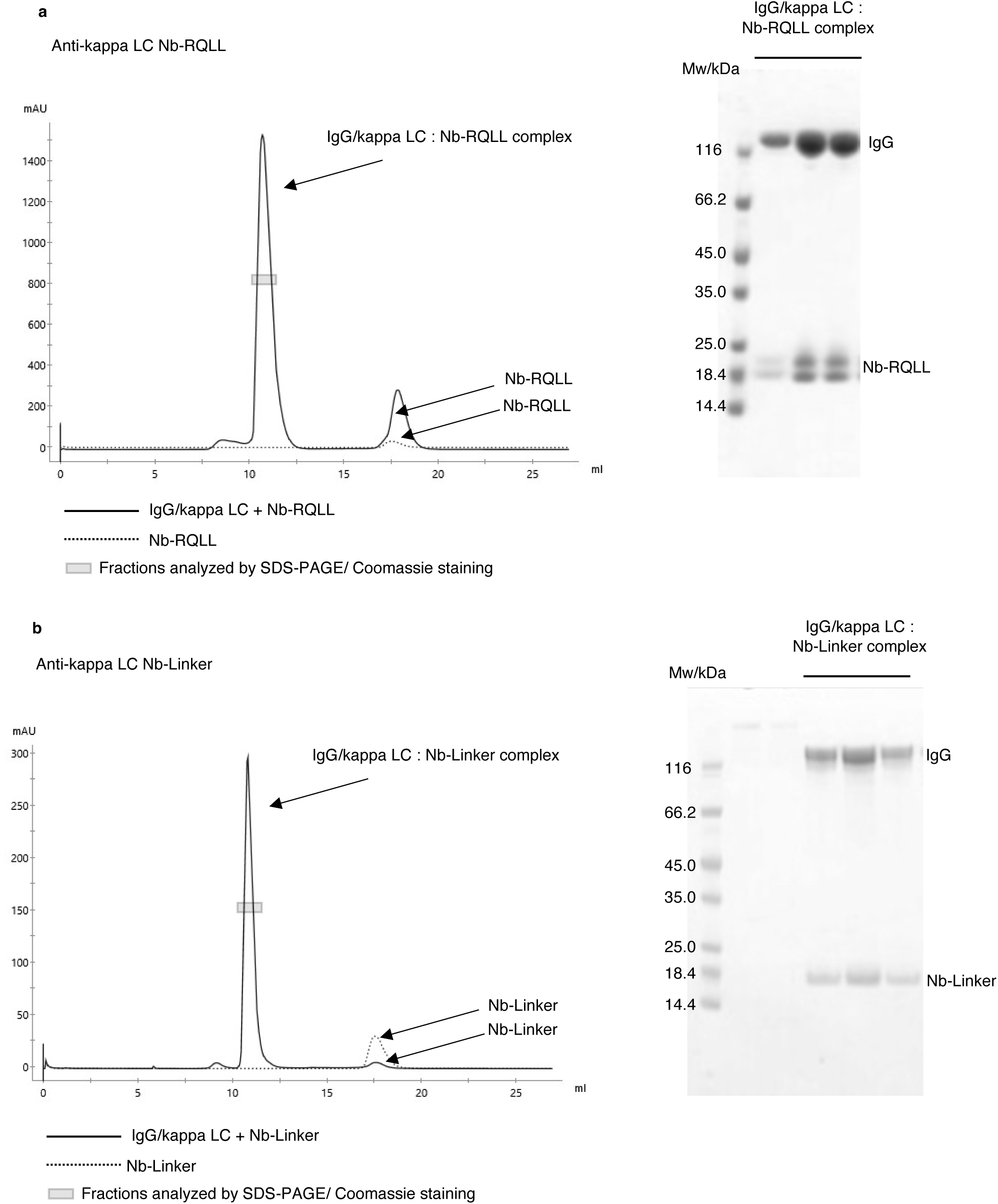
**a-b,** Size exclusion chromatography confirming complex formation between IgG containing kappa LC and Nb-RQLL (**a**) and Nb-Linker (**b**). Non-reducing SDS-PAGE (Commassie staining) showing co-elution of IgG with Nb-RQLL **(a**) or Nb-Linker (**b**), respectively, in peak fractions as indicated with box in chromatogram.

**Extended data Fig. 4.**
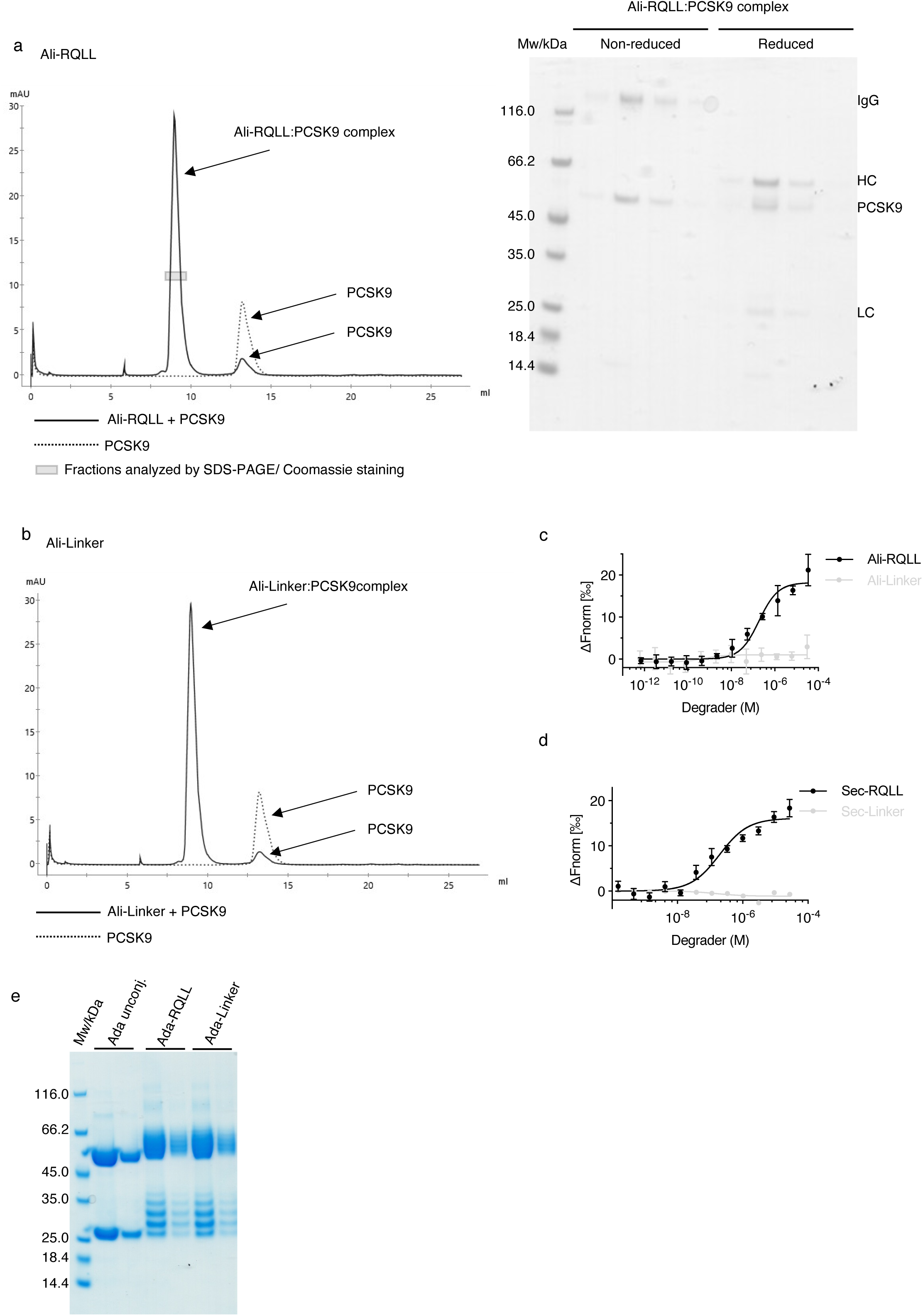
**a,** Gel filtration chromatogram (right) showing co-elution of PCSK9 and Ali-RQLL. SDS-PAGE (Commassie staining) of peak fractions (indicated by box in chromatogram) containing PCSK9 and Ali-RQLL. Intact IgG (non-reducing conditions) and heavy chains (HC)/light chains (LC) (reducing conditions) are indicated. **b,** Gel filtration showing co-elution of PCSK9 and Ali-Linker. **c,** The binding of Ali-RQLL and Ali-linker to sortilin ECD was analyzed using MST. The titration of antibody ranged from 34.2 µM to 0.7 pM (Ali-RQLL) or 30.6 µM to 0.6 pM (Ali-Linker) with a constant concentration of fluorescently labelled sortilin ECD at 100 nM. The change in thermophoretic signal allowed calculation of the kD = 189 ± 67 nM for the interaction between Ali-RQLL and sortilin ECD. In contrast, no interaction could be measured between Ali-Linker and Sortilin (n=4). **d,** The binding of Sec-RQLL and Sec-linker to sortilin ECD was analyzed using MST. The titration of antibody ranged from 27 µM to 155 pM (Sec-RQLL) or 28 µM to 157 pM (Sec-Linker) with a constant concentration of fluorescently labelled sortilin at 100 nM. The change in thermophoretic signal allowed calculation of the kD = 185 ± 73 nM for the interaction between Sec-RQLL and sortilin, while no interaction could be detected between Sec-Linker and sortilin. Error bars represent standard deviation (n=4). e, Linkage of RQLL peptide to antibodies resulted in a mass shift of approximately 8-12 kDa in molecular weight of the HC and LC fragments as visualized by reducing SDS-PAGE. This indicates that each antibody was linked at 2-10 positions. n indicates the number of biological replicates in each experiment. Each data set shown is representative of at least three independent experiments.

**Extended data Fig. 5.**
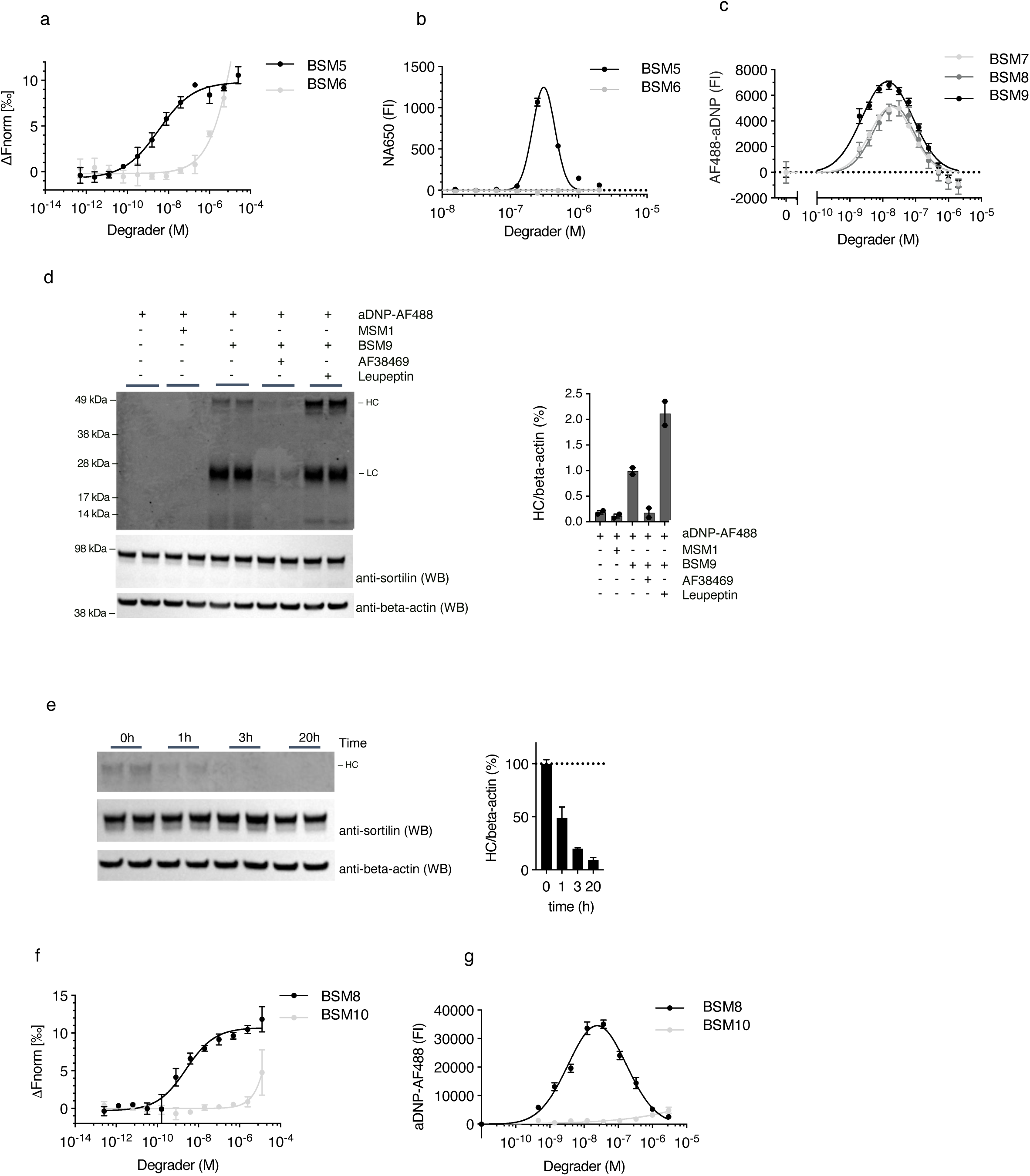
**a-b,** SORTAC activity dependents on sortilin binding as replacing the sortilin binding part with an inactive enantiomer (BSM6) resulted in complete loss of affinity for sortilin ECD as measured by MST (mean ± SD, n=2) (**a**) as well as cellular uptake of NA650 (100 nM) by HEK293/sortilin cells (3 h incubation) (Data are shown as mean ± SD, n=2) compared to the corresponding active SORTACs (BSM5) (**b**). **c,** Three DNP-SORTACs with varying linker lengths (BSM7-9) induced dose-dependent cellular uptake of aDNP-AF488 IgG (100 nM) in HEK293/sortilin cells (3 h incubation). Data shown are shown as mean ± SEM (n=2). **d**, Fluorescent Western blot of HEK293/sortilin cells lysates incubated for 6 h with aDNP-AF488 (100 nM), SORTAC BSM9 (30 nM), leupeptin (80 µM), and sortilin monobinders MSM1 (30 nM) and AF38469 (10 µM) as indicated. A band corresponding to IgG heavy chain (HC) was detected in lysates from cells co-incubated with target and BSM9 degrader. The intensity of the band was markedly increased upon inhibition of lysosomal degradation by leupeptin, and absent in cells where the sortilin monobinders (AF38469) or MSM1 was added. The two lower gels are Western blots of the same samples for sortilin and beta-actin, respectively. Bar graph shows quantification of HC band FI signal normalized to beta-actin signal (mean±SEM) (n=2). **e,** Time course experiment showing the disappearance of internalized aDNP-AF488 following SORTAC-induced internalization in HEK293/sortilin cells, analyzed by fluorescent Western blotting of lysates of HEK293/sortilin cells. Cells were incubated for 3 h with aDNP-AF488 (100 nM) and BSM9 (30 nM) where after the cells were washed and the culture supernatant was exchanged with fresh medium without target and SORTAC. Cells were subsequently lysed at the indicated time points. The bar graph shows quantification of the HC band normalized to the beta-actin band (Data are shown as mean ± SEM) (n=2). **f-g**, Exchange of active sortilin binder in BSM10 with its inactive enantiomer in BSM8, resulted in loss of binding to sortilin ECD in MST (mean ± SD, n=2) (**f**) and inability to induced internatilization of aDNP-AF488 in HEK293/sortilin cells (3 h incubation) (mean value ± SEM, n=2) (**g**), showing the critical requirement of intact sortilin binding. n indicates the number of biological replicates in each experiment. Each data set shown is representative of at least three independent experiments.

**Extended data Fig. 6.**
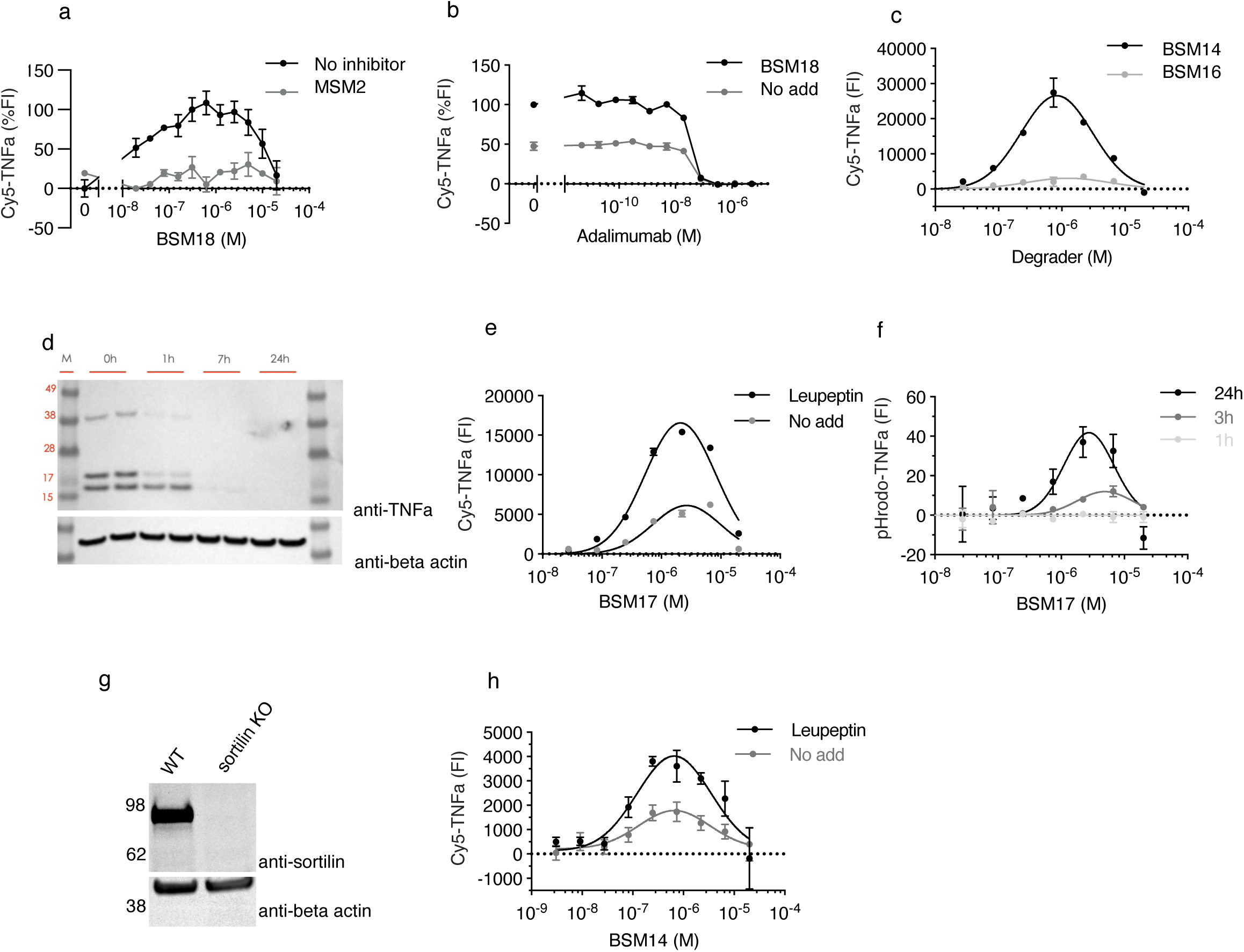
**a,** SORTAC-induced Cy5-TNFa (100 nM) uptake in HEK293/sortilin cells (24h) was inhibited by the presence of excess i sortilin monobinder (MSM2, 20 µM) (mean value ± SEM, n=2). **b**, Cy5-TNFa uptake (100 nM) induced by SORTAC BSM18 (1 µM) in HEK293/sortilin cells was inhibited by increasing concentration of adalimumab. Data shown as mean value ± SD (n=2). **c,** a SORTAC based on the inactive enantiomer (BSM16) of the sortilin binder did not induce cellular uptake of TNFa (100 nM) in HEK293/sortilin cells. In contrast, the active enantiomer SORTAC BSM14 induced potent dose-dependent uptake. Data shown as mean ± SEM (n=2) **d,** Western blot for TNFa showing that internalized TNFa induced by SORTAC (BSM17, 3 µM) over the course of 3 h in HEK293/sortilin cells appears as three distinct bands that rapidly disappear over time following removal of TNFa and SORTAC from the culture supernatant (upper panel). Western blot for beta-actin is included as loading control (lower panel). **e,** SORTAC-induced cellular uptake during 24 h of Cy5-TNFa (100 nM) in HEK293/sortilin cells in the presence or absence of the lysosomal proteinase inhibitor leupeptin (80 µM), showing that the Cy5 signal markedly increased in the presence of leupeptin. Data are shown as mean ± SEM (n=2). **f,** Concentration and time-dependent SORTAC-induced cellular uptake of TNFa (100 nM) labeled with the pH-sensitive fluorophore pHrodo (pHrodo-TNFa) by HEK293/sortilin cells. Data are shown as mean ± SD (n=2). **g,** Western blot analysis of Schwann cell lysates from WT and sortilin KO rats, respectively, using the indicated antibodies. **h,** SORTAC-induced internalization of Cy5-TNFa (100 nM) in WT Schwann cells in the presence or absence of leupeptin (80 µM). Data are shown as mean ± SEM, n=2). n indicates the number of biological replicates in each experiment. Each data set shown is representative of at least three independent experiments.

